# The diversification of mealybugs was triggered by new symbiont acquisitions and followed by adaptive radiations on host plants

**DOI:** 10.1101/2024.10.23.619971

**Authors:** Jinyeong Choi, Pradeep Palanichamy, Seunghwan Lee, Filip Husnik

## Abstract

Symbiotic microorganisms play a critical role in supplementing beneficial nutrients to herbivorous insects feeding on unbalanced diets. These microbial symbionts can both facilitate or constrain plant-feeding insects’ adaptations to certain host plants, depending on their gene content and metabolic potential. The diet breadth of herbivorous insects is considered an important evolutionary factor affecting genotypic and phenotypic changes associated with host shifts. Acquiring new symbionts can, therefore, drive changes in niche breadth and subsequent adaptive radiation(s). Mealybugs comprise one of the major groups of scale insects, most of which feed on diverse angiosperms. Different sub-lineages of mealybugs also house different lineages of bacteria and fungi as their obligate symbionts. Here, we use mealybugs as a model system to test the hypothesis that the evolution of herbivorous insects is driven by both obligate symbionts and host plants. Based on metagenome analyses of 28 host species as well as a literature survey, we identified Betaproteobacteria, Gammaproteobacteria, Flavobacteriia, and *Ophiocordyceps* fungi as obligate symbionts of the major clades of mealybugs. A time-calibrated phylogenetic tree of mealybugs allowed us to infer the ancestral obligate symbionts of the major mealybug clades. Our results indicate that the emergence of major mealybug lineages coincided with the acquisitions of new obligate endosymbionts. Subsequent radiations of mealybugs were inferred to have mostly resulted from the adaptive radiation through continuous host shifts on angiosperms. The contribution of microbial symbiosis to the diversification of herbivorous insects is thus likely limited by new symbiont origins or replacements, and insect adaptations play a larger role in further plant switches.

## Introduction

Half of all described insect species (over 400,000) are herbivorous, largely due to the abundance and ubiquity of plants on Earth (Strong et al., 1984; Grimaldi & Engel, 2005). Understanding the ecological factors driving the remarkable diversification of both insects and plants remains a key evolutionary question. The diversification of herbivorous insects is thought to be driven by shifts to new host plants or adaptations to different ecological niches of their original hosts (Futuyama & Agrawal, 2009). However, the role of microbial symbionts in the evolution of herbivorous insects is increasingly being recognized.

Plant sap-feeding insects have mutualistic relationships with diverse microbial symbionts (McCutcheon & Moran, 2012). These symbionts, which can be either obligatory or facultative for their host, play nutritional and defensive roles, such as essential amino acid biosynthesis or protecting against plant toxins and parasitoids (Hammer & Bowers, 2015; Douglas, 2016; Vorburger, 2021). The dietary and ecological niches of herbivorous insects are greatly influenced by their symbiotic microbes, as these microbes can either enhance or restrict their use of host plants (Hansen & Moran, 2014; Bennett & Moran, 2015). The acquisitions of new symbionts have been proposed to broaden the dietary options of insects and drive their evolutionary diversification (Janson et al., 2008; Bennett & Moran, 2015; Sudakaran et al., 2015; Sudakaran et al., 2017). The effect of microbial symbionts is strikingly evident in the species-rich order Hemiptera (over 82,000 species), which has a long history of fluctuating associations with symbiotic microbes (Sudakaran et al., 2017).

Mealybugs are a prominent group of scale insects, with approximately 2,300 species, most of which primarily feed on angiosperms (García Morales et al., 2024). Because some species are polyphagous, they are considered economic pests, damaging diverse plants through sap-feeding and virus transmission (McKenzie, 1967; Kondo et al., 2008; Tsai et al., 2010; García Morales et al., 2024). Mealybugs include three families: Pseudococcidae, Rhizoecidae, and Xenococcidae (Choi & Lee, 2022). The Pseudococcidae family contains three subfamilies, Rastrococcinae, Phenacoccinae, and Pseudococcinae (Choi & Lee, 2022). While many species of Phenacoccinae and Pseudococcinae are cosmopolitan, Rastrococcinae primarily inhabit subtropical or tropical regions (Williams, 2004). The Rhizoecidae and Xenococcidae primarily feed on plant roots, and they are found in all continental regions (Kozár & Konczné Benedicty, 2007).

Mealybugs have distinct obligate endosymbionts that seem to largely mirror their main clades, including flavobacterial symbionts in Rastrococcinae, Rhizoecidae, and Xenococcidae; Betaproteobacteria in Phenacoccinae; and Betaproteobacteria + Gammaproteobacteria in Pseudococcinae (Choi & Lee, 2022). Of these symbionts, the betaproteobacterial genus *Candidatus* Tremblaya has gained attention for its unique features. Notably, the Pseudococcinae mealybugs have the “bacteriumwithin-bacterium” symbiotic system, with gammaproteobacterial symbionts living within *Tremblaya princeps* (von Dohlen et al., 2001). These symbionts collaborate to synthesize essential amino acids and vitamins (McCutcheon & von Dohlen, 2011; Husnik et al., 2013; Husnik & McCutcheon, 2016; Garber et al., 2021). On the other hand, a single symbiont, *Tremblaya phenacola*, has been identified in the sister subfamily Phenacoccinae (Gruwell et al., 2010; López-Madrigal et al., 2014). Interestingly, genome sequencing revealed that *Tremblaya phenacola* from *Phenacoccus peruvianus* Granara de Willink and *P. solenopsis* Tinsley contains a chimeric genome of mixed betaproteobacterial and gammaproteobacterial origin (Gil et al., 2018; Bai et al., 2024).

Despite the progress in mealybug symbiont genome evolution, the evolutionary history of mealybugs with their symbiotic microbes and host plants remains poorly understood. We hypothesize that new acquisitions of nutritional symbionts occurred early during mealybug evolution, leading to the emergence of different sub-lineages specialized to distinct ecological niches. To test this hypothesis here, we: (i) screened microbial symbionts of mealybugs using metagenome sequencing and a literature review, (ii) performed a time-calibrated phylogenetic analysis of mealybugs using available fossil data, (iii) conducted ancestral state reconstruction for obligate symbionts on the deep mealybug nodes, and (iv) characterized the feeding preferences of major mealybug lineages.

## Materials & Methods

### DNA extraction and metagenome screening of endosymbionts

A total of 28 species of mealybugs belonging to four major lineages (Phenacoccinae, Pseudococcinae, Rastrococcinae and Rhizoecidae) were selected for metagenome sequencing (Table S1). The samples were preserved in 99% ethanol at -20°C. For DNA extraction, 5–22 individuals of each species were selected, and the wax secretions were removed using a pair of fine forceps under a dissecting microscope. Any individuals showing parasitoid symptoms were excluded at this step. The samples were then thoroughly washed with 99% ethanol to reduce surface contaminants. Genomic DNA was extracted using the MasterPure Complete DNA Purification Kit (Epicenter, USA) according to the manufacturer’s protocols. The extracted DNA was quantified using a Qubit 4 fluorometer (Thermo Fisher Scientific, Waltham, MA, USA). DNA library preparation and sequencing were carried out using the Illumina NovaSeq 6000 sequencer with 2 × 150 bp paired-end reads. The quality of the raw Illumina reads was assessed using FastQC v0.11.7 (Andrews, 2010), and low-quality reads and adapters were filtered with fastp (v.0.20.0) (Chen et al., 2018) if necessary. The approximate taxonomic affiliation and relative abundance of microbes were determined from 16S and 18S rRNA read abundance in metagenomic data using phyloFlash v3.4 (Gruber-Vodicka et al., 2020). Additional endosymbionts of mealybugs were further identified from various literature sources (Table S2).

### Molecular phylogenetic analyses of endosymbionts

The raw Illumina reads were assembled *de novo* using SPAdes v3.15 (Prjibelski et al., 2020) with default parameters or the following combination of k-mers (21, 33, 55, 77, 99, 111, 119 and 127). The rRNA sequences of microbes showing a high abundance in phyloFlash were extracted using barrnap v3 (https://github.com/tseemann/barrnap) and identified through BLASTn searches. Any other rRNA sequences of symbionts from mealybugs, other hosts, and free-living microbes, were retrieved from NCBI (accession numbers in Figure S1). The 16S rRNA and 23S rRNA sequences for bacteria and 18S-28S rRNA (including ITS and 5.8S) sequences for fungi were aligned using the MAFFT v7 (*--auto*) (Katoh & Standley, 2013). The 16S rRNA and 23S rRNA alignments were concatenated with Sequencematrix v1.7.8 (Vaidya et al., 2011). Maximum likelihood (ML) analyses were performed with the two datasets (bacteria and fungi) using IQ-tree v1.6.12 (Nguyen et al., 2015). The best-fitting models were selected by ModelFinder (Kalyaanamoorthy et al., 2017) as implemented in IQ-tree (TVMe+I+G4 for bacterial 16S + 23S rRNA alignments and TIM3+F+G4 for fungal 18S-28S rRNA alignments). Branch support was assessed through ultrafast bootstrap (UFBoot) approximation with 1,000 replicates (Hoang et al., 2018). The ML trees were visualized in Figtree v1.4.4.

### Sequence matrix and alignment for host insects

A total of 110 species, including 96 ingroup species of Rhizoecidae, Xenococcidae and Pseudococcidae, and 14 outgroup species of Coccidae, Diaspididae, and Kermesidae, were included in the phylogenetic analyses of host insects. We selected sequences of eight loci (*COI, 18S, 28S D2* and *D10, Dynamin, Ef-1α 5’* and *3’* and *wingless*) that were previously used in mealybug phylogenetic analyses (Choi & Lee, 2022). The sequences for each species were obtained from the assemblies generated in this study or from NCBI (Table S3). Using Geneious Prime v2022.1.1 (Kearse et al., 2012), the reference sequences of each locus were mapped to the assembled contigs of each metagenome assembly, and the consensus sequences were confirmed through BLAST searches to ensure they were derived from the mealybug genomes. The acquired sequences were aligned using MAFFT v7 (-auto) and edited following the procedures outlined in Choi & Lee (2022), which involves deleting introns from *Dynamin, Ef-1α 5’* and *3’* sequences, translating protein-coding sequences of *COI, Dynamin, Ef-1α 5’* and *3’*, and *wingless* into amino acids to check for stop codons, trimming ambiguous and poorly aligned sequences with Gblocks 0.91b (Castresana, 2000; Talavera & Castresana, 2007) using relaxed parameters, and combining the edited sequences of each locus using Sequencematrix v1.7.8. Information on symbionts was available for *Mirococcus clarus, Peliococcus calluneti, Phenacoccus avenae, Rhodania porifera*, and *Vryburgia brevicruris*, but they were not included in this phylogenetic analysis due to the lack of available molecular data of the hosts.

### Divergence time estimation

The divergence times were estimated using BEAST v2.6.2 (Bouckaert et al., 2019) with a relaxed clock model (Drummond et al., 2006) and Bayesian inference via Markov chain Monte Carlo (MCMC). A Yule model was used as the prior for species tree estimation (Yule, 1925). The MCMC chain was run for 10 million generations, with samples collected every 10,000 generations. The stationarity, convergence, and effective sample sizes (ESS) of the analysis were evaluated using Tracer v1.7.1 (Rambaut et al., 2018). The GTR model was applied uniformly to the concatenated sequence data. A maximum clade credibility tree was constructed using TreeAnnotator v2.6.2 (Bouckaert et al., 2019) after discarding 25% of the trees as burn-in. The tree was visualized using Figtree v1.4.2 (Rambaut, 2009). We used four fossil calibrations including Coccidae (*Rosahendersonia prisca* from Burmese amber), Diaspididae (*Normarkicoccus cambayae* from Cambay amber), Kermesidae (*Sucinikermes kulickae* from Baltic amber), and Pseudococcidae (*Williamsicoccus megalops* from Lebanese amber) to date the divergence times, corresponding to the oldest fossil records of each group (Vea & Grimaldi, 2016). The fossil species *W. megalops* was used as the Pseudococcidae clade representative, including the Rastrococcinae, Phenacoccinae, and Pseudococcinae based on its morphological features. This species was described based on an alate male exhibiting two pairs of glandular pouches on abdominal segments VII and VIII, which is a characteristic known from the Phenacoccinae and Rastrococcinae, but not present in the Rhizoecidae and Xenococcidae (Vea & Grimaldi, 2015; Hodgson, 2020). The most recent common ancestors were set to 104 ± 6 Ma for Coccidae, 75 ± 25 Ma for Diaspididae, 72.5 ± 27.5 Ma for Kermesidae, and 137.5 ± 2.5 Ma for Pseudococcidae (Vea & Grimaldi, 2015; Vea & Grimaldi, 2016). Each group was constrained to be monophyletic according to the current classification and molecular phylogenies (Hodgson & Hardy, 2013; Choi & Lee, 2022).

### Ancestral state reconstruction

To infer the evolutionary history of symbiont acquisitions and replacements in mealybugs, we estimated the ancestral character states of selected nodes on the host tree using BayesTraits v3.0.1 (Pagel, 2004; Pagel and Meade, 2006). Based on our symbiont screening results, six lineages of microorganisms were identified as obligate symbionts of mealybugs as follows: *Candidatus* Brownia rhizoecola (Flavobacteriia) in Rhizoecidae and Xenococcidae, two lineages of Flavobacteriia and two lineages of *Ophiocordyceps* in Rastrococcinae, and *Tremblaya* (Betaproteobacteria) in Phenacoccinae and Pseudococcinae. To estimate the posterior probabilities and values of these six traits at ancestral nodes of the host phylogeny, we used reversible-jump Markov chain Monte Carlo (MCMC) methods in BayesTraits, with the BayesMultiState model. The rate deviation was set to 10, and the MCMC chains were run for 10 million generations, sampling parameters every 1000th generation, and discarding the first one million generations as burn-in. Although Gammaproteobacteria are almost always detected in Pseudococcinae species as co-obligate symbionts of *Tremblaya*, they were excluded from this analysis for simplicity as they are additionally established symbionts that have been acquired multiple times (Thao et al., 2002; Husnik and McCutcheon 2016).

### Characterizing the feeding preference of mealybugs

We compiled the host-plant information for 1,784 species in the Pseudococcidae clade, 135 species in the clade Rhizoecidae + Xenococcidae, and 25 species in the Rastrococcinae clade. The data were sourced from ScaleNet (García Morales et al., 2024; accessed September 2020). The feeding patterns of mealybugs were characterized by the host-plant range and type. The host-plant range was presented as mean values with standard deviations for each group, based on the number of recorded host plant families and genera per species. Further, the herbivores were defined as monophagous (feeding on a single plant genus), oligophagous (feeding on two or more genera within a family), and polyphagous (feeding on two or more families) (Cates, 1980). The host-plant type was displayed as the composition ratio of angiosperms, gymnosperms, and pteridophytes. We indicated “present” in the respective column for each plant category if a species had an occurrence record on gymnosperms, pteridophytes, and angiosperms. Multiple recorded occurrences of a single species on different host plants, including gymnosperms, pteridophytes, and angiosperms, were treated as separate instances. Therefore, the total number of host records (marked as “present”) exceeded the number of examined species.

## Results

### Diversity and phylogenetic relationships of mealybug endosymbionts

The microbial endosymbionts of mealybugs were identified as bacteria belonging to the classes Flavobacteriia, Alphaproteobacteria, Betaproteobacteria, and Gammaproteobacteria, and the fungal genus *Ophiocordyceps* based on the phylogenetic analyses of rRNA sequences (Figure 1). This result revealed that most microbes from metagenomes of 28 mealybug species (with asterisks in Figure S1) were closely related to previously reported symbionts in mealybugs and other insects except for a few symbionts that cluster at distinct phylogenetic positions. Rastrococcinae were found to harbor symbionts from Flavobacteriia, Alphaproteobacteria, and Gammaproteobacteria, as well as *Ophiocordyceps* (Figures 1 and S1). *Rastrococcus iceryoides* and the other *Rastrococcus* species harbor distinct lineages of flavobacterial symbionts. The flavobacterial symbiont of *R. iceryoides* was determined to be closely related to *Shikimatogenerans* of bostrichid beetles, while the clade including the flavobacterial symbiont of the other *Rastrococcus* species was sister to a clade with symbionts from other scale insect families. The *Rastrococcus* symbionts, excluding the *R. iceryoides* symbiont, were tentatively considered a single lineage, although they are paraphyletic due to the inclusion of *Skilesia* and the symbiont of *Cryptococcus ulmi*, which requires confirmation through further sampling. *R. mangiferae* and *R. tropicasiaticus* were found to house different strains of *Ophiocordyceps*, related to those found in soft scales. Additionally, *Rastrococcus* species contained symbionts from the genera *Arsenophonus, Rickettsia, Wolbachia*, and another Gammaproteobacteria.

**Figure 1.**
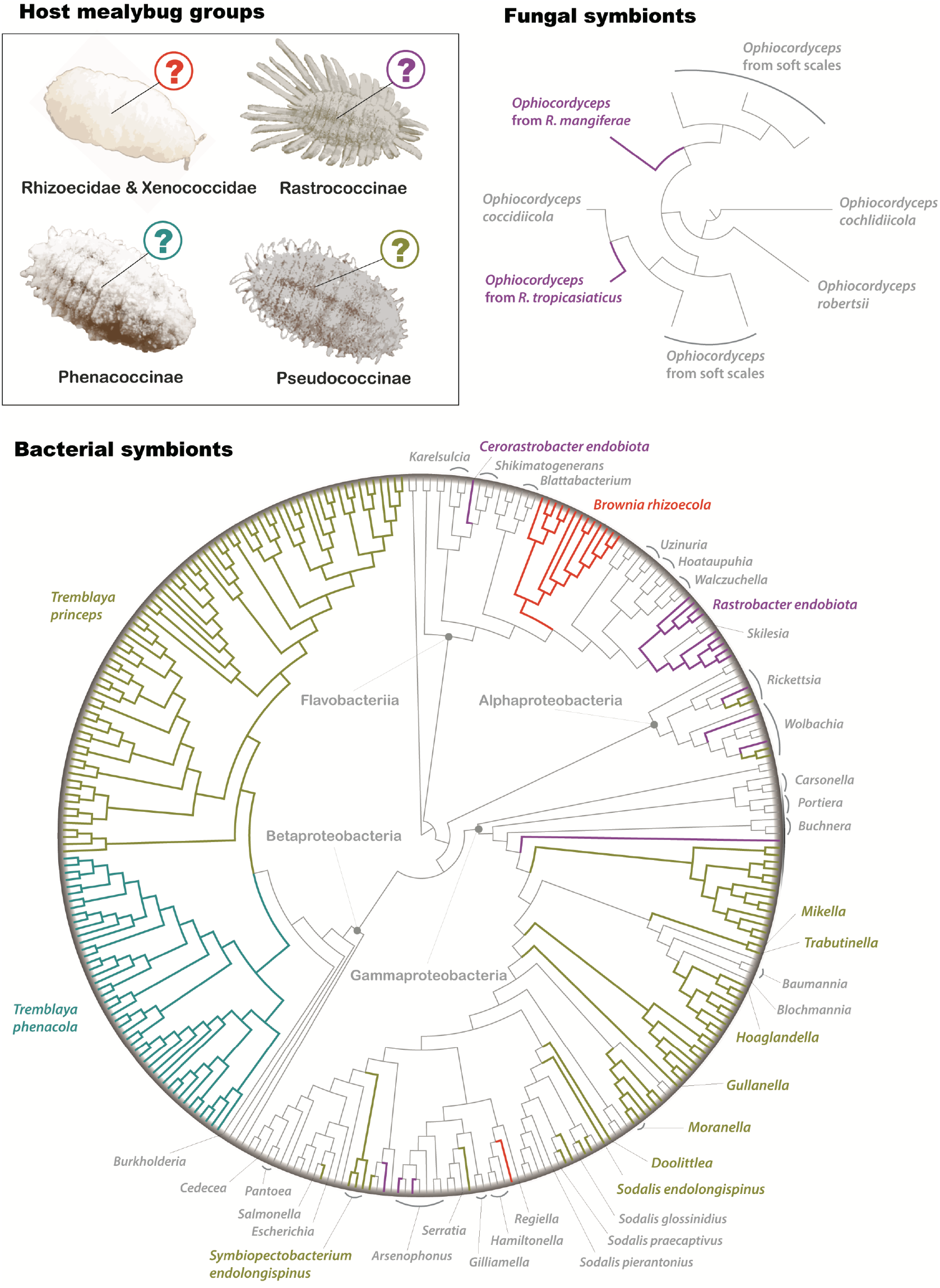
Phylogenetic trees of mealybug symbionts. The trees were inferred with maximum likelihood (ML) in IQ-tree based on 5,542 bp (for bacterial tree) and 1,942 bp (for fungal tree) of sequences from the rRNA region. Symbionts of mealybugs are highlighted with different colors according to their insect hosts. The other branches in gray represent symbionts of other insects and free-living microbes. Detailed information on terminal taxa and node values is available in Figure S1.

Rhizoecidae harbor *Brownia rhizoecola* (Flavobacteriia) and Gammaproteobacteria (Figures 1 and S1). *Brownia rhizoecola* formed a clade (UFBoot = 100) that was sister to a clade including male-killing endosymbionts of ladybugs, *Skilesia* of aphids, and symbionts from other scale insect families. *Rhizoecus amorphophalli* was found to also harbor a gammaproteobacterial symbiont related to *Hamiltonella defensa* and *Regiella insecticola* from aphids.

Phenacoccinae and Pseudococcinae primarily contain *Tremblaya* (Betaproteobacteria) and diverse symbionts within Alphaproteobacteria and Gammaproteobacteria (Figures 1 and S1). *Tremblaya* formed a clade (UFBoot = 100), sister to a free-living *Burkholderia thailandensis*. This clade of *Tremblaya* included two subclades of *T. phenacola* (UFBoot = 92) and *T. princes* (UFBoot = 100). *T. phenacola* was solely detected in Phenacoccinae species that do not contain intrabacterial co-symbionts, while Pseudococcinae species showed the presence of one to four additional symbionts from Alphaproteobacteria and Gammaproteobacteria, as well as *T. princeps*. These cosymbionts represent multiple lineages related to *Rickettsia, Serratia, Sodali, Wolbachia*, and gammaproteobacterial symbionts of other insects, as well as symbionts of the scale insect genus *Puto*.

### Phylogeny and divergence times of mealybugs

The time-estimated phylogenetic tree showed distinct clades for the major mealybug lineages: Rhizoecidae + Xenococcidae, Rastrococcinae, Phenacoccinae, and Pseudococcinae (Figures 2 and S2). The most recent common ancestor (MRCA) of these groups was estimated to have originated approximately 141 million years ago (Ma) [135.8–147.6 Ma based on 95% highest posterior densities]. Subsequently, the MRCA of the Rastrococcinae, Phenacoccinae, and Pseudococcinae was estimated to 137 Ma [134.7–139.5 Ma]. The MRCA of Phenacoccinae and Pseudococcinae was estimated to 125 Ma [119.7–130.6 Ma]. This divergence time of the Phenacoccinae + Pseudococcinae clade is younger than the estimate (135–140 Ma) in Vea & Grimaldi (2016). Even though we used the same fossil calibrations, our study additionally includes Rastrococcinae that are likely responsible for the difference in dating. The MRCA of Rhizoecidae and Xenococcidae was estimated to have originated at 109 Ma [94.1–123.8 Ma]. The MRCAs of each family and subfamily were estimated to have originated in the mid to late Cretaceous. The MRCA of Rhizoecidae was estimated at 84 Ma [76.0–92.7 MA]. The MRCAs of each Phenacoccinae and Pseudococcinae were estimated at 105 Ma [97.6–112.5 Ma] and 108 Ma [101.6–117.4 Ma], respectively. The MRCA of Rastrococcinae was estimated to have appeared at 72 Ma [63.8–81.1 Ma]. In short, the origin of the three major deep nodes of mealybugs was estimated to have occurred in the early Cretaceous. Subsequently, the species diversification of each mealybug lineage began in the mid-Cretaceous, with the subsequent radiations accelerating during the Cenozoic era.

**Figure 2.**
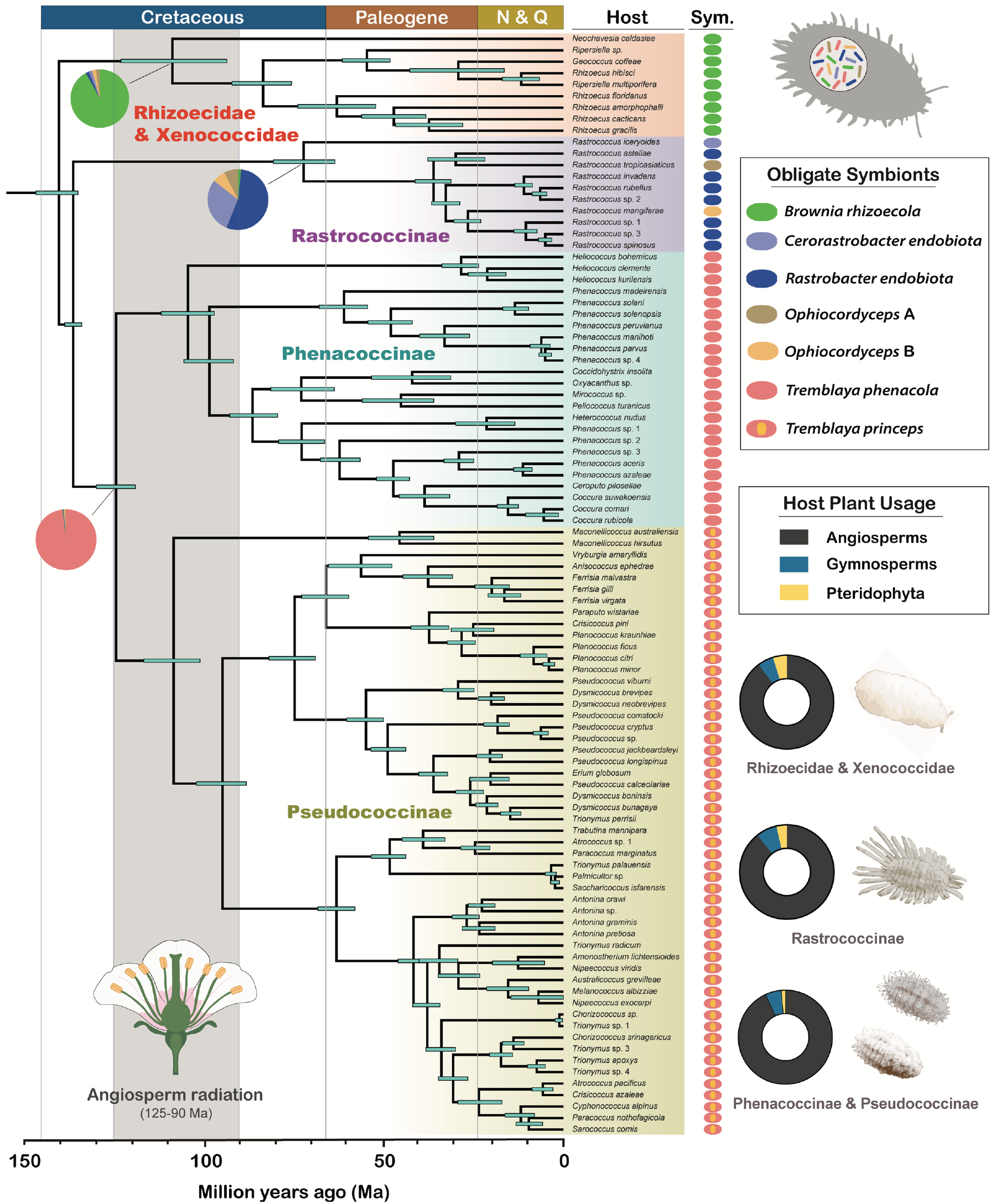
Time-calibrated phylogeny of mealybugs and ancestral state reconstruction for their obligate symbionts. Divergence times were estimated by BEAST under a relaxed-clock model with four fossil calibrations. The maximum clade credibility tree was constructed with TreeAnnotator. Bars on each node represent the 95% highest posterior density (HPD) interval. The median posterior estimates and posterior probabilities are available in Figure S2. Ancestral state reconstruction was performed with Bayestraits. Pie charts represent percent probabilities of ancestral obligate endosymbionts. Obligate symbionts of each species of mealybug are shown on the right-hand side of species labels. The host plant usage of each mealybug lineage is represented with pie charts (see detailed information in Table 4).

### Ancestral symbiont state reconstruction

The origin of obligate symbiotic associations was estimated for three key nodes of the mealybug phylogenetic tree (Figure 2; pie charts). The main outcomes of each node are summarized as follows: (i) *Tremblaya* had the highest reconstructed probability (> 98%) at the node representing the MRCA of Phenacoccinae + Pseudococcinae. Low probabilities of the other symbionts were inferred in this node from 0.01% to 0.05. (ii) *Brownia rhizoecola* had the highest reconstructed probability (> 92%) at the node representing the MRCA of Rhizoecidae + Xenococcidae. The other symbionts showed low probabilities for this node from 0.4% to 2.2%. (iii) *Rastrobacter endobiota* had the highest reconstructed probability (55%) at the node representing the MRCA of Rastrococcinae, followed by *Cerorastrobacter endobiota* (29%) and the other symbionts with much lower probabilities from 0.9% to 7.3%.

### Feeding preferences of major mealybug lineages

The host-plant types were analyzed by the number of host records for each mealybug group (Figure 2; Table S4). In the case of Pseudococcidae, the largest number of records was from angiosperms (1,710 records; 93.09%), with the remaining proportion comprising gymnosperms (101 records; 5.50%) and pteridophytes (26 records; 1.42%). Rhizoecidae and Xenococcidae showed the largest number of host records from angiosperms (131 records; 89.73%), with the remaining proportion comprising gymnosperms (8 records; 5.48%) and pteridophytes (7 records; 4.79%). For Rastrococcinae, the largest number of records was also from angiosperms (25 records; 89.29%), with the remaining proportion comprising gymnosperms (2 records; 7.14%) and pteridophytes (1 record; 3.57%). The host plant ranges for the major lineages of mealybugs were quantified at both the plant familyand genus level (Table S4). At the family level, the mean host-plant ranges were calculated as follows: 3.7 ± 5.9 for Rhizoecidae + Xenococcidae, 5.5 ± 8.1 for Rastrococcinae, and 2.8 ± 6.9 for Pseudococcidae. At the genus-level, the mean hostplant ranges were calculated as follows: 5.2 ± 9.3 for Rhizoecidae + Xenococcidae, 8.7 ± 17.7 for Rastrococ-cinae, and 4.9 ± 16.3 for Pseudococcidae. Among the examined mealybug species, 56% were monophagous, followed by 32% polyphagous and 12% oligophagous.

### “*Candidatus* Cerorastrobacter endobiota” and “*Candidatus* Rastrobacter endobiota”

Among bacterial symbionts, one of the flavobacterial lineages was previously considered to be an obligate endosymbiont of *Rastrococcus* (Gruwell et al., 2010; Choi & Lee, 2022). In this study, we found an additional flavobacterial lineage from *Rastrococcus iceryoides*, which has a distinct phylogenetic position from the known flavobacterial symbiont of other *Rastrococcus*. We propose the designations “*Candidatus* Cerorastrobacter endobiota” for the flavobacterial symbiont of *R. iceryoides* and “*Candidatus* Rastrobacter endobiota” for the flavobacterial symbionts of the other *Rastrococcus* species. “*Rastro-*” denotes the genus of the associated hosts, while the suffix “*-bacter*” indicates their bacterial affiliation. The prefix “*Cero-*” in *Cerorastrobacter* reflects the host’ trait of heavy wax secretion. The term “*endobiota*”, derived from the Latin “*endo*” meaning “within” and “*biota*” referring to the living organisms, signifies the presence of these bacteria as endosymbionts residing within the host insects.

## Discussion

### Ancestral obligate endosymbionts of major mealybug lineages

Several new clades of bacterial and fungal symbionts were identified from the Rastrococcinae in this study (Figures 1 and 2). The topology of *Rastrobacter endobiota* was mostly congruent with that of its host insects (Figures 2 and S1), which suggests codivergence between some hosts and symbionts. Furthermore, it was closely related to *Skilesia* of the aphid genus *Geopemphigus* and other scale insect families (Figures 1 and S1), indicating their free-living common ancestor independently established symbioses with scale insects and aphids. On the other hand, *Cerorastrobacter endobiota* has a long branch and is clustering with *Candidatus* Karelsulcia and *Candidatus* Shikimatogenerans, which are ancient symbionts of auchenorrhynchan insects and bostrichid beetles (Moran et al., 2005; McCutcheon & Moran, 2007; Kiefer et al., 2023). Among the fungal symbionts, two distinct strains of *Ophiocordyceps* were found from *R. mangiferae* and *R. tropicasiaticus*. Buchner (1965) reported the existence of symbiotic yeasts in some *Rastrococcus* species. Although he also examined the yeasts in *R. iceryoides*, no distinct fungal contigs were identified in its metagenomes in this study. To date, no genome sequencing, transmission electron microscopy (TEM), or fluorescence *in situ* hybridization (FISH) studies have been performed on the symbionts of Rastrococcinae, so their roles and localization within the hosts remain largely unknown. Nevertheless, based on the available evidence, we conclude that *Candidatus* Cerorastrobacter endobiota, *Candidatus* Rastrobacter endobiota, and two *Ophiocordyceps* are very likely obligate endosymbionts of Rastrococcinae.

*Brownia rhizoecola* was a predominant symbiont found in the Rhizoecidae and Xenococcidae (Figures 1 and S1). Gruwell et al. (2010) proposed *B. rhizoecola* to be an obligate endosymbiont of these root mealybugs. Our phylogenetic analysis suggests that *B. rhizoecola* is monophyletic, although the taxon sampling is still relatively limited. The topology of *B. rhizoecola* was not completely congruent with that of its host insects at the species level, but a similar deep branching pattern was observed for several subclades (Figures 2 and S1). Further phylogenomic analyses for the hosts and symbionts are thus needed to robustly resolve the evolutionary history of *Brownia* in root mealybugs. *B. rhizoecola* was found to be closely related to symbionts of various insects such as cockroaches, auger beetles, ladybugs, aphids, and other scale insects (Figures 1 and S1). We cannot rule out phylogenetic artifacts affecting this topology, but it appears that their common free-living ancestor was capable of colonizing diverse insect hosts and evolved into beneficial symbionts in some of them. The localization of *B. rhizoecola* within the host bacteriocytes was confirmed using TEM and FISH (Szklarzewicz et al., 2022). However, the exact role of *B. rhizoecola* in the host remains unknown due to the absence of genomic data. Based on the available evidence, our data suggest that *B. rhizoecola* is an ancestral obligate endosymbiont of both Rhizoecidae and Xenococcidae.

*Tremblaya phenacola* and *T. princeps* were uniformly identified from every Phenacoccinae and Pseudococcinae species we used for sequencing (Figures 1 and S1). That *Tremblaya* is an obligate endosymbiont of these mealybug subfamilies is well known (Thao et al., 2002; Gruwell et al., 2010) and our data confirm these results. The phylogenetic analysis revealed that *Tremblaya* is monophyletic, and most branches were congruent with those of their host insects (Figures 2 and S1). *T. phenacola* and *T. princeps* were found to be sister clades, confirming their common ancestry and co-diversification with mealybug hosts. Furthermore, *Tremblaya* was most closely related to *Burkholderia* (Figures 1 and S1), implying that it evolved from a free-living bacterium into a nutritional symbiont (Urban & Cryan, 2012). The genomes of *T. phenacola* and *T. princeps* are highly reduced but they retain genes for the biosynthesis of essential amino acids (Husnik & McCutcheon, 2016). Both *Tremblaya princeps* and *phenacola* were observed to reside within host bacteriocytes through TEM and FISH (von Dohlen et al., 2001; Koga et al., 2013; López-Madrigal et al., 2014; Garber et al., 2021). All available data support that *Tremblaya* is an ancestral obligate endosymbiont of Phenacoccinae and Pseudococcinae.

### Independent acquisitions and replacements shape the mealybug endosymbioses

Our ancestral state reconstruction revealed that three major establishments of microbial symbionts initially occurred in the major mealybug lineages during the Cretaceous period (Figure 3). The common ancestor of the Phenacoccinae and Pseudococcinae acquired *Tremblaya* at approximately 125 Ma. Following the diversification of their insect hosts, *Tremblaya* diverged into two different lineages, *T. phenacola* and *T. princeps*. Additionally, multiple gammaproteobacterial strains were introduced to the Pseudococcinae species, becoming co-obligate endosymbionts residing within *T. princeps. Brownia* was predicted to be acquired by the common ancestor of Rhizoecidae and Xenococcidae at 109 Ma. After the diversification, some root mealybugs likely acquired Gammaproteobacteria as a co-symbiont, as evidenced by *Rhizoecus* species harboring an additional lineage of symbionts related to *Hamiltonella* and *Regiella* (Figure 1). Although the aforementioned host lineages have retained *Brownia* and *Tremblaya* as their main obligate symbionts, the Rastrococcinae have undergone symbiont replacements during their diversification. *Rastrobacter* was predicted to be established in the common ancestor of Rastrococcinae at 73 Ma. However, some species in this group have lost this symbiont and acquired new bacterial or fungal symbionts, such as *Cerorastrobacter* and *Ophiocordyceps*. Additionally, alphaproteobacterial and gammaproteobacterial symbionts were introduced to the Rastrococcinae species. The fluctuation of obligate nutritional symbionts within this single host clade indicates that Rastrococcinae potentially alternate their diet between complete and incomplete nutritional composition. Many scale insects can likely feed on plant cell contents, which provide a more nutritionally balanced diet than phloem sap that lacks essential amino acids (Gullan & Kosztarab, 1997).

**Figure 3.**
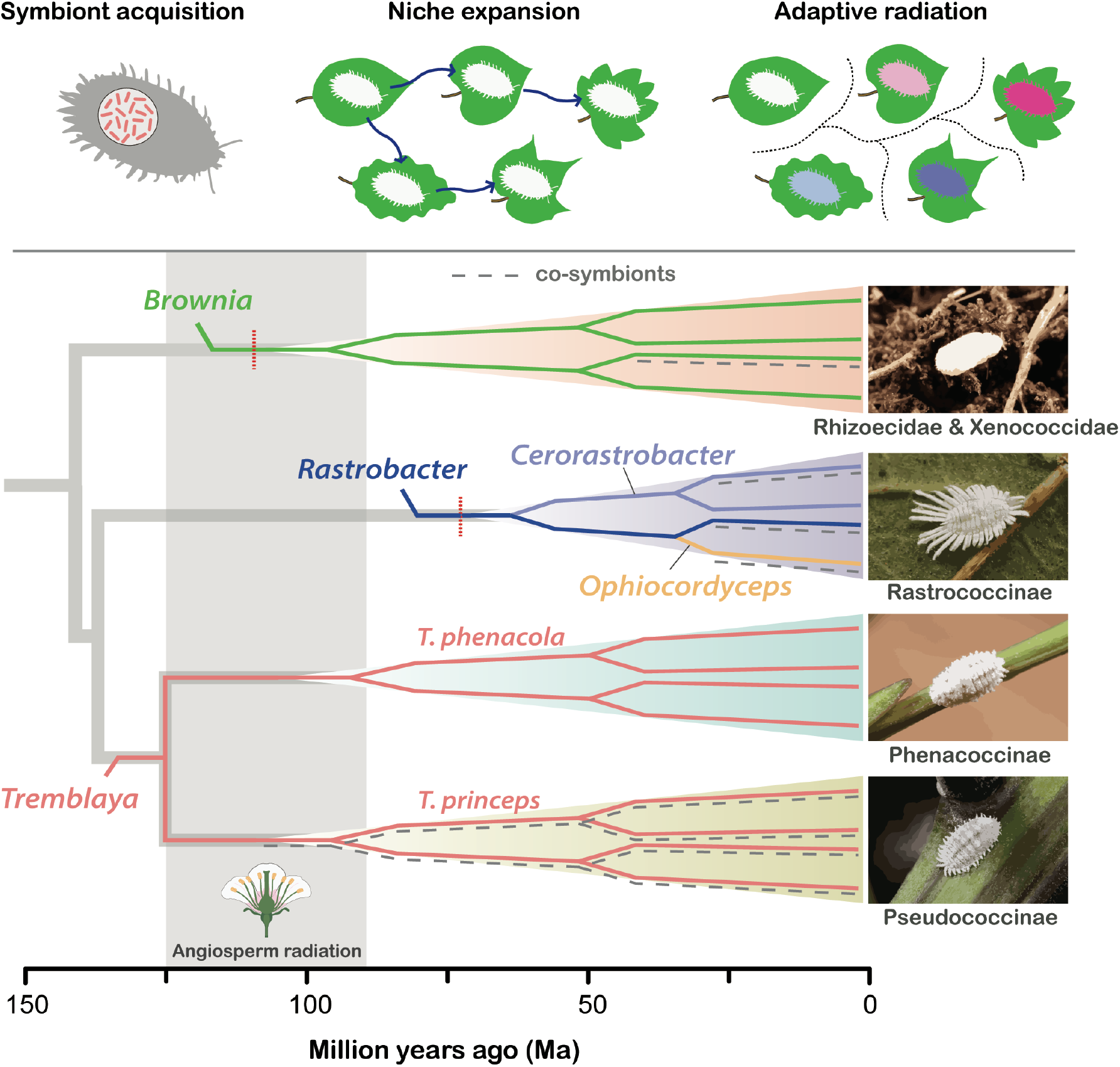
Schematic diagram of hypothetical scenarios of the mealybug diversification associated with symbiont replacements and angiosperm radiation. Solid color lines represent obligate symbionts of each mealybug group. Co-symbionts (when present) are shown as gray dash lines under the lines of obligate symbionts.

### Mealybugs exploit diverse angiosperms with considerable variation in host-plant range

Although all major clades of mealybugs primarily use angiosperms as their host plants (Table S4), some mealybugs feed on conifers, cycads, lichens, ferns, and other gymnosperms and pteridophytes. This plant host pattern is typical for scale insects, with over 97% of extant species known to feed on angiosperms, with some exceptions (e.g. Matsucoccidae) (Gullan & Cook, 2007; Kozár, 2004; Vea & Grimaldi, 2016). Our study revealed significant variation in diet breadth among mealybug species (Table S4). Like other plant-feeding insects that tend to have highly specialized and conservative feeding modes (Forister et al., 2015), most mealybug species are also monophagous or oligophagous (Table S4). However, over 30% of mealybugs are polyphagous, with broad host ranges. Among them, *Ferrisia virgata* (Cockerell), *Maconellicoccus hirsutus* (Green), *Planococcus citri* (Risso), *Pseudococcus viburni* (Signoret), and *Phenacoccus solenopsis* Tinsley are notoriously polyphagous with host records of over 200 plant genera (García Morales et al., 2024). The diet breadth of herbivores can be influenced by environmental and ecological factors, including latitude and the plant species richness in their habitat (Agosta et al., 2010; Forister et al., 2015). In mealybugs, dispersal constraints due to reduced legs and occasional passive wind dispersal likely affected the host range expansion in some species (Normark, 2011). Scale insects are atypically displaying broader host use at lower latitudes, which was proposed to result from their passive dispersal (Hardy et al., 2015). The host specialization is potentially costly for scale insects if not coevolving in parallel with mechanisms that increase the chance of finding the host plant. On the other hand, the evolution of polyphagy allows random settlements on neighboring plants and more efficient feeding. This contrasts with closely related groups, such as aphids, which typically possess a winged stage capable of long-distance dispersal and allowing more efficient localization of specific host plants (Dixon, 1977). A potential downside of polyphagy is that it increases the exposure to the chemical defense of various plants. How polyphagous mealybugs overcome these defenses is unfortunately still very poorly understood. Alongside host plant specialization, host range expansion emerges as a preferred evolutionary strategy for some mealybugs as it allows them to thrive in environments with a high diversity of flowering plants, such as tropics and subtropics.

### Diversification of mealybugs

The ancestor of each major mealybug lineage acquired its obligate endosymbiont during the Cretaceous (Figure 3) which was characterized by the radiation and diversification of angiosperms (Crane et al., 1995). The subsequent diversification of mealybugs appears to have accelerated in the Late Cretaceous and Cenozoic, following the rapid radiation of angiosperms. A similar pattern has been observed in most insect herbivores (Peris & Condamine 2024), especially in the Pyrrhocoridae, where the transition of core symbionts coincided with the radiation of their main host plants, Malvales, and preceded the major diversification of the pyrrhocorid bugs (Sudakaran et al., 2015). The acquisition of novel symbionts can allow their hosts to access new ecological niches (Cornwallis et al., 2023). In addition, the host switch of plant-feeding insects to preexisting, closely related host plants can increase the rate of their diversification (Percy et al., 2004). Some mealybugs that successfully colonized new plant taxa may experience adaptive radiations and divergent selection-driven speciation as they become subject to different selection pressures. This divergence can be facilitated by the physical and chemical diversity of angiosperm hosts, which increases the heterogeneity of insect environments (Futuyma & Agrawal, 2009). Although monophagous and oligophagous species make up a large proportion of mealybugs, the role of polyphagous feeding in diversification cannot be ruled out. The oscillation hypothesis suggests that a broader diet breadth in herbivores can provide the foundation for host plant-associated speciation, while also facilitating geographical range expansion (Janz & Nylin, 2008; Slove & Janz, 2011). Therefore, polyphagy in some mealybugs likely contributed to the expansion of their populations into new ecological niches, promoting specialization and speciation. Overall, our results suggest that the emergence of major mealybug lineages was likely triggered by the acquisition of new obligate endosymbionts. However, the subsequent mealybug radiations were unlikely driven by novel symbionts but rather resulted from sympatric or allopatric speciation through host shifts on angiosperms.

## Acknowledgments

JYC was supported by the National Research Foundation of Korea (NRF) grant funded by the Korean government (MSIT) (2021R1A6A3A03038909) and the JSPS KAKENHI grant (20939772). FH was supported by the JSPS KAKENHI grant (23K14256) and the HFSP Early Career Grant (RGEC29/2024). In addition, this work was supported by a grant from the National Institute of Biological Resources (NIBR), funded by the Ministry of Environment (MOE) of the Republic of Korea (NIBR202406201). We acknowledge Dr Hirotaka Tanaka (Ehime Univ.) and Dr Thibaut Malausa (INRAE, France) for providing us samples for this study. We thank Penny Gullan (Australian National Univ.) for giving us feedback on this manuscript. We also thank the Scientific Computing (SCDA) and Sequencing (SQC) sections of the Okinawa Institute of Science and Technology for their great support.

---

**Figure S1a.**
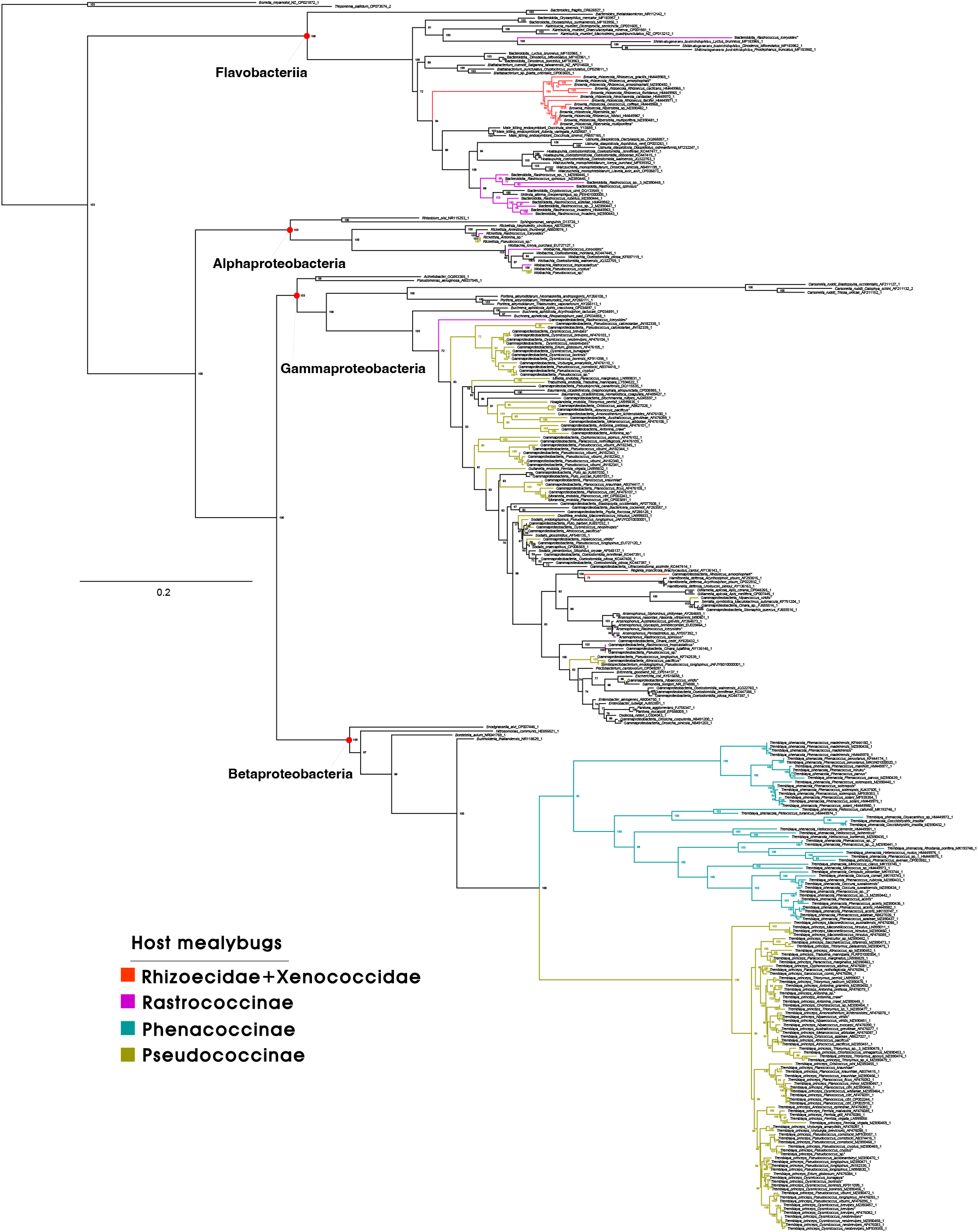
Maximum likelihood (ML) tree of bacterial symbionts of mealybugs, inferred from 5,542 bp of 16S rRNA and 23S rRNA sequences in IQ-tree. The numbers at each node represent ultrafast bootstraps. Tip labels consist of symbiont identities, host insects (absent in freeliving bacteria), and Genbank accession numbers. Asterisks (*) indicate samples extracted from metagenomes in this study. Symbionts of mealybugs color-coded according to their insect hosts.

**Figure S1b.**
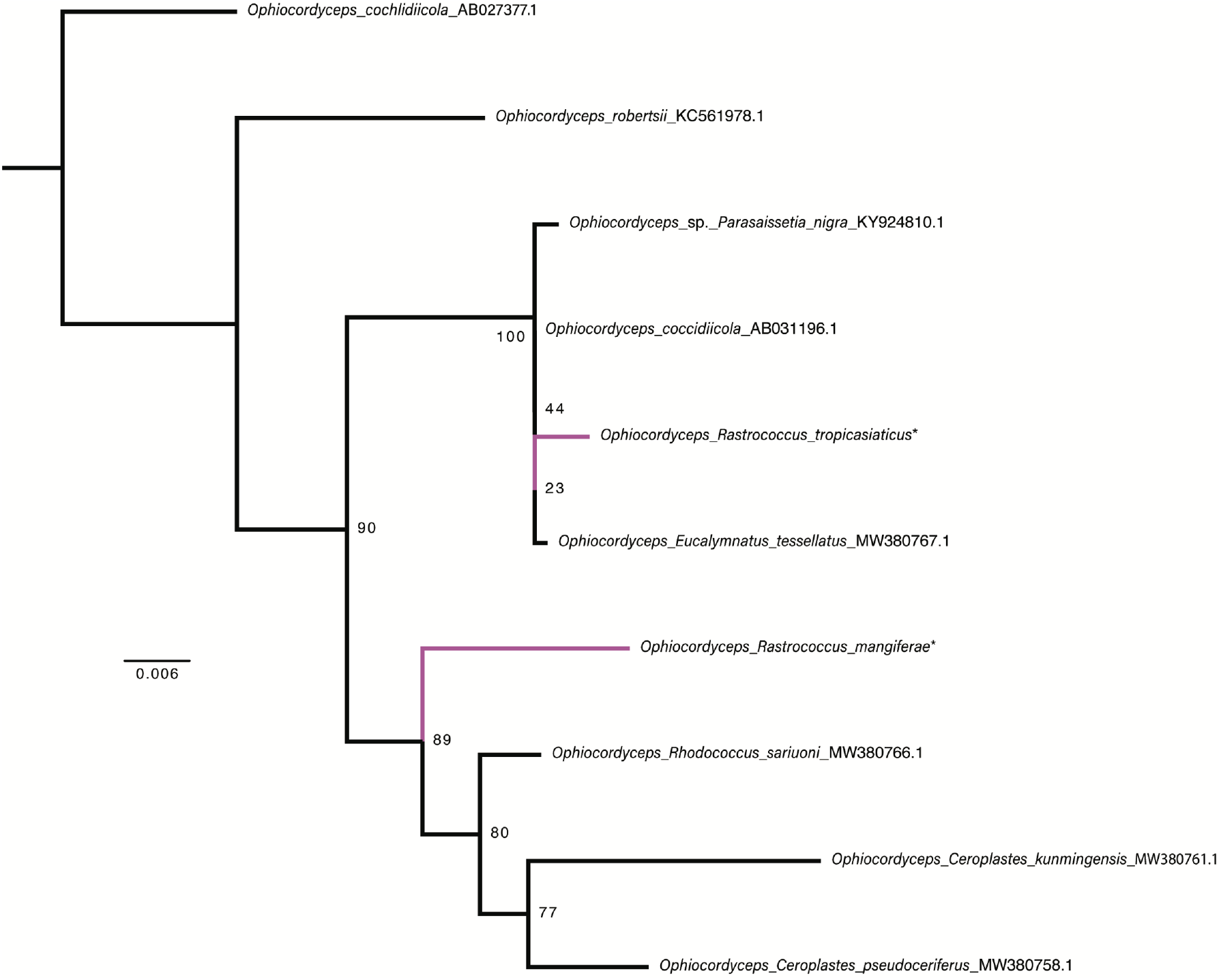
Maximum likelihood (ML) tree of fungal symbionts of mealybugs, inferred from 1,942 bp of 18S–23S rRNA sequences in IQ-tree. The numbers at each node represent ultrafast bootstraps. Tip labels consist of symbiont identities, host insects (absent in free-living fungi), and Genbank accession numbers. Asterisks (*) indicate samples extracted from metagenomes in this study. Symbionts of mealybugs are highlighted in purple.

**Table S1.**
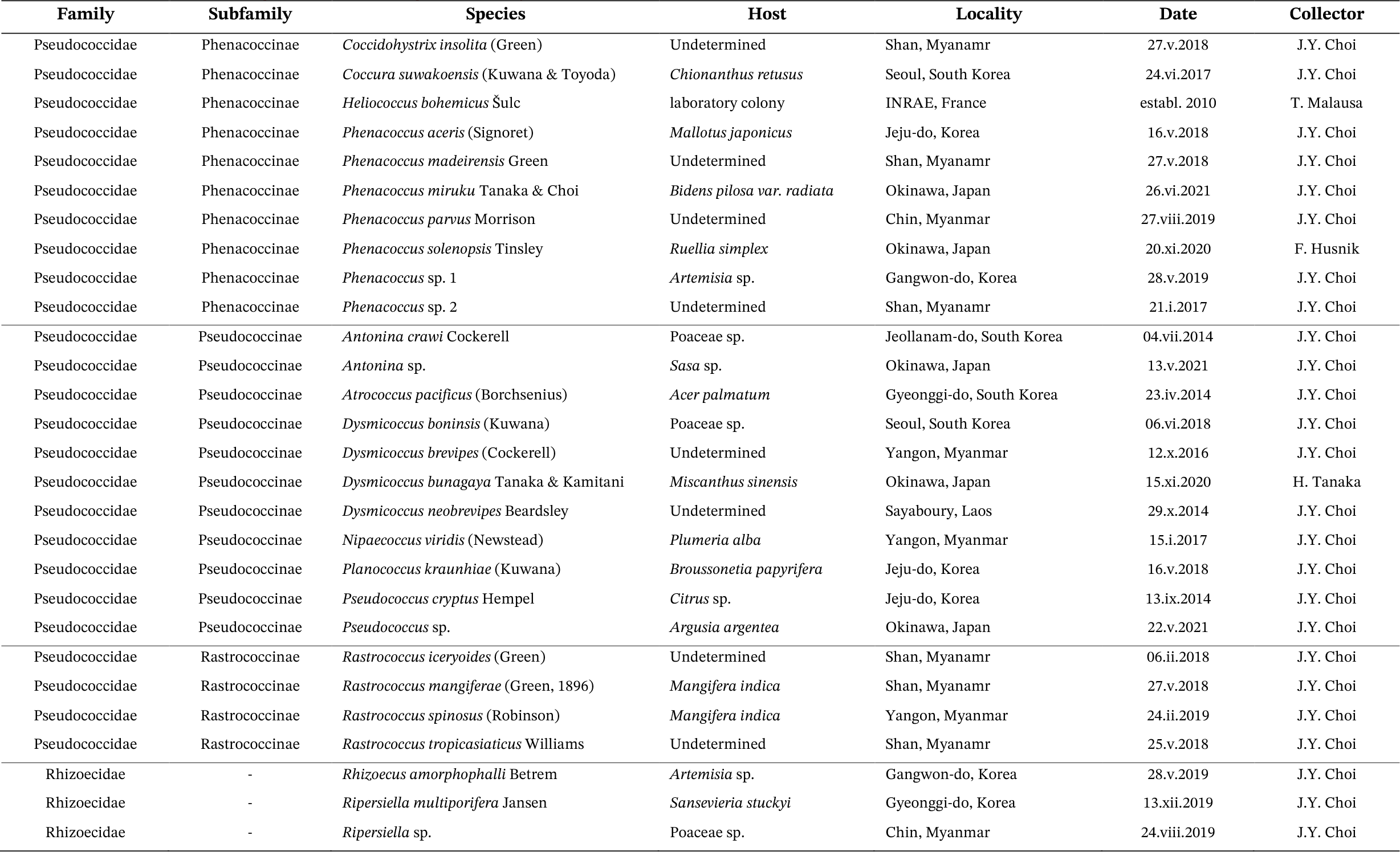
Samples used for metagenome sequencing in this study.

**Table S2.**
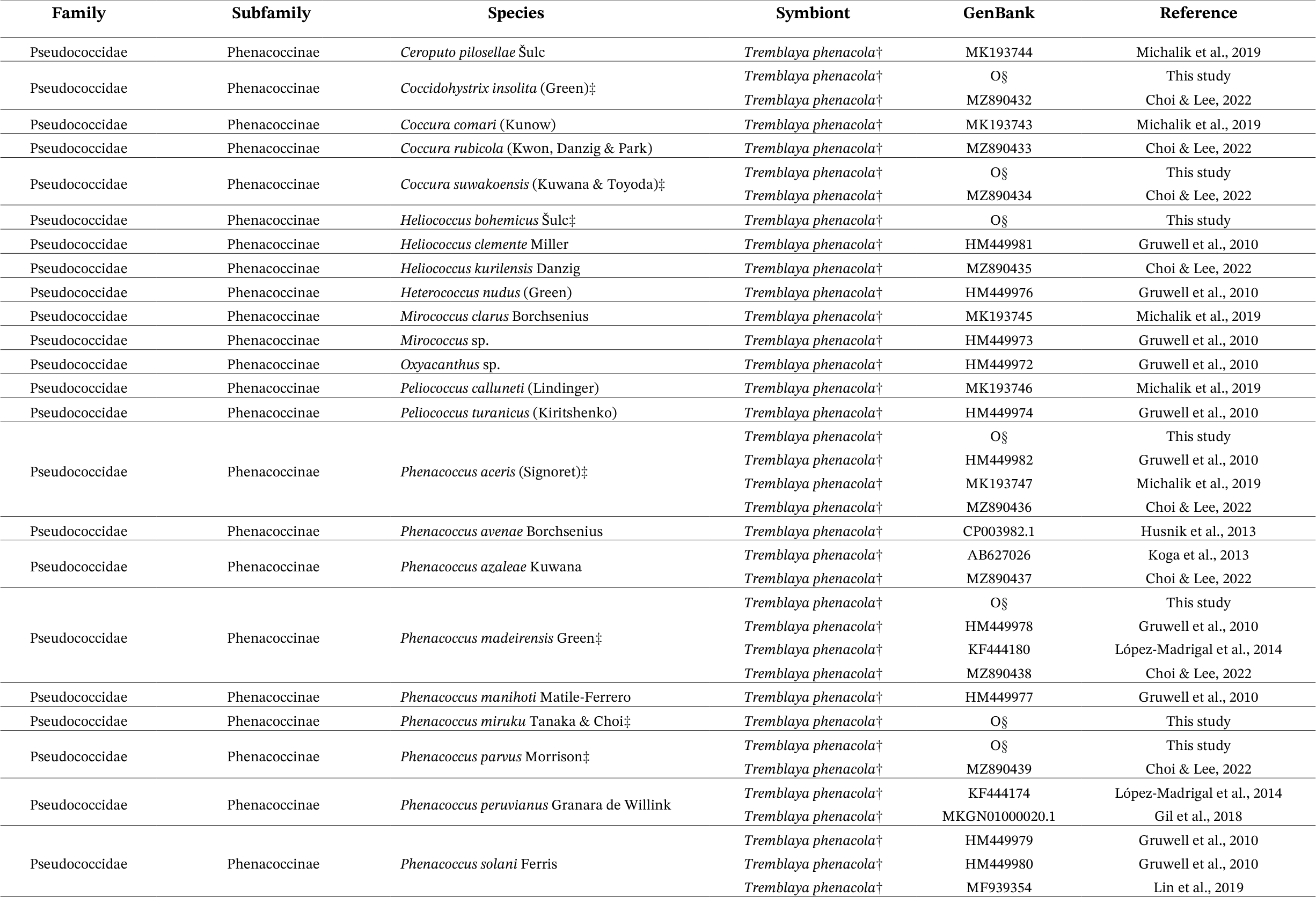
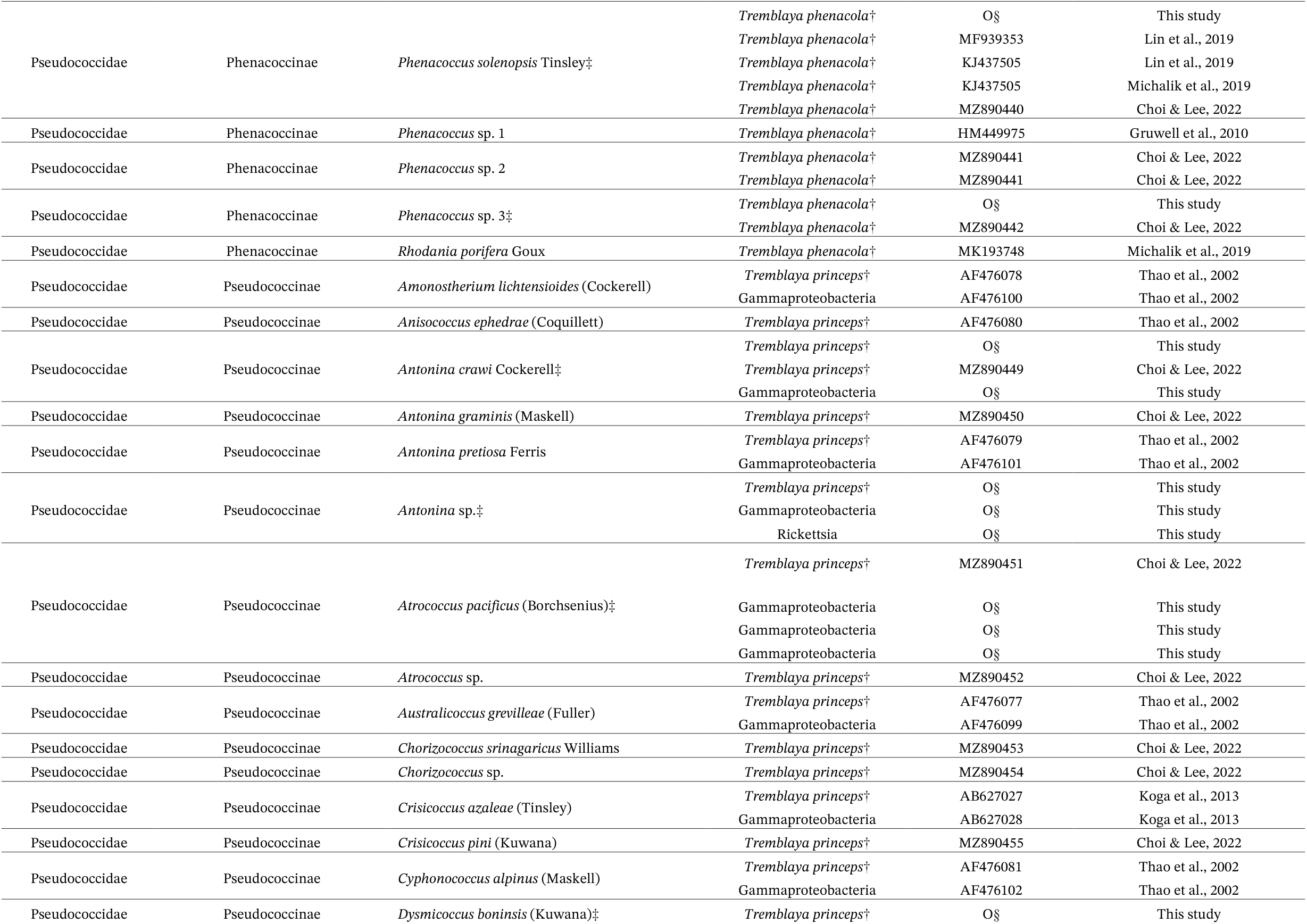
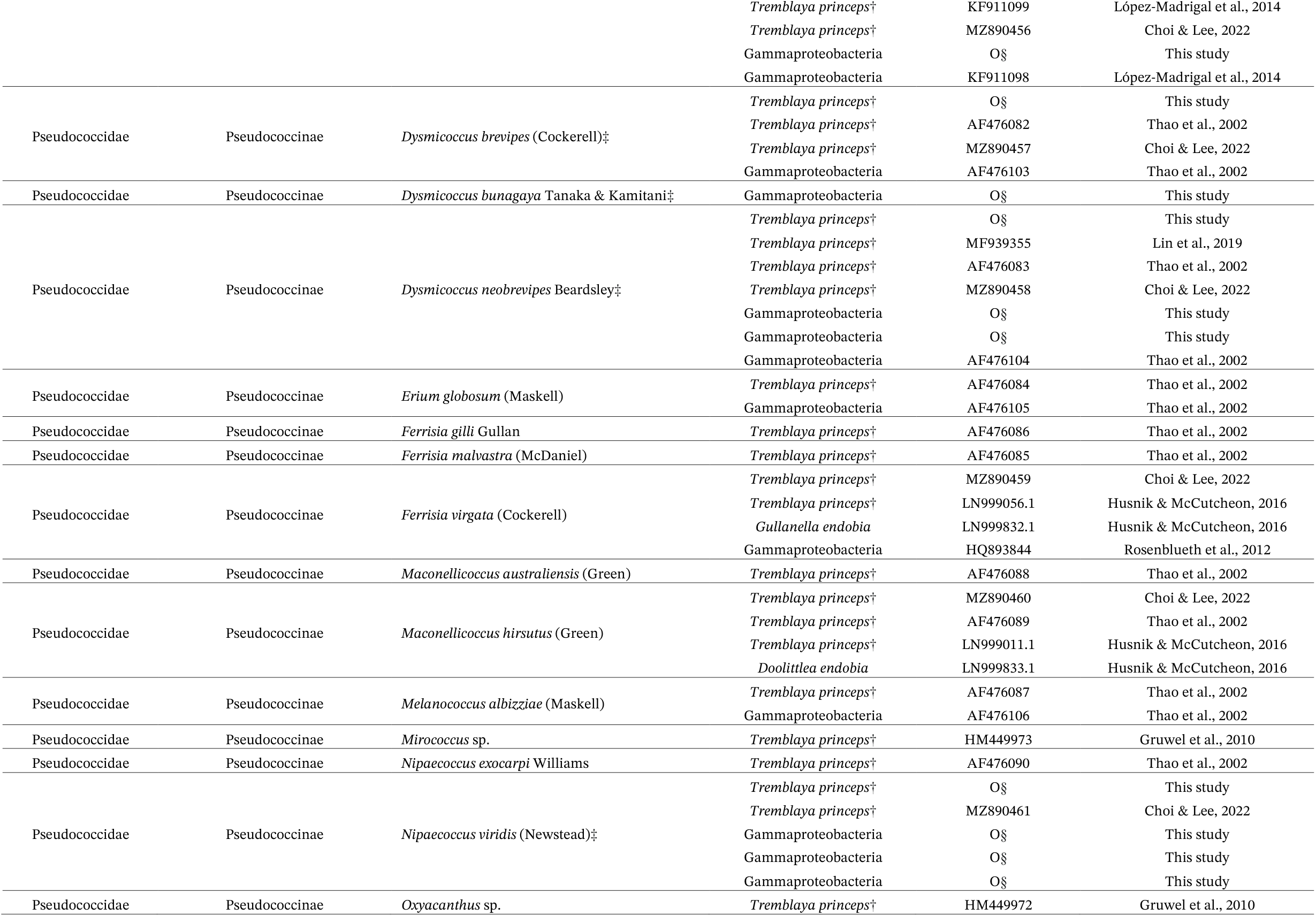
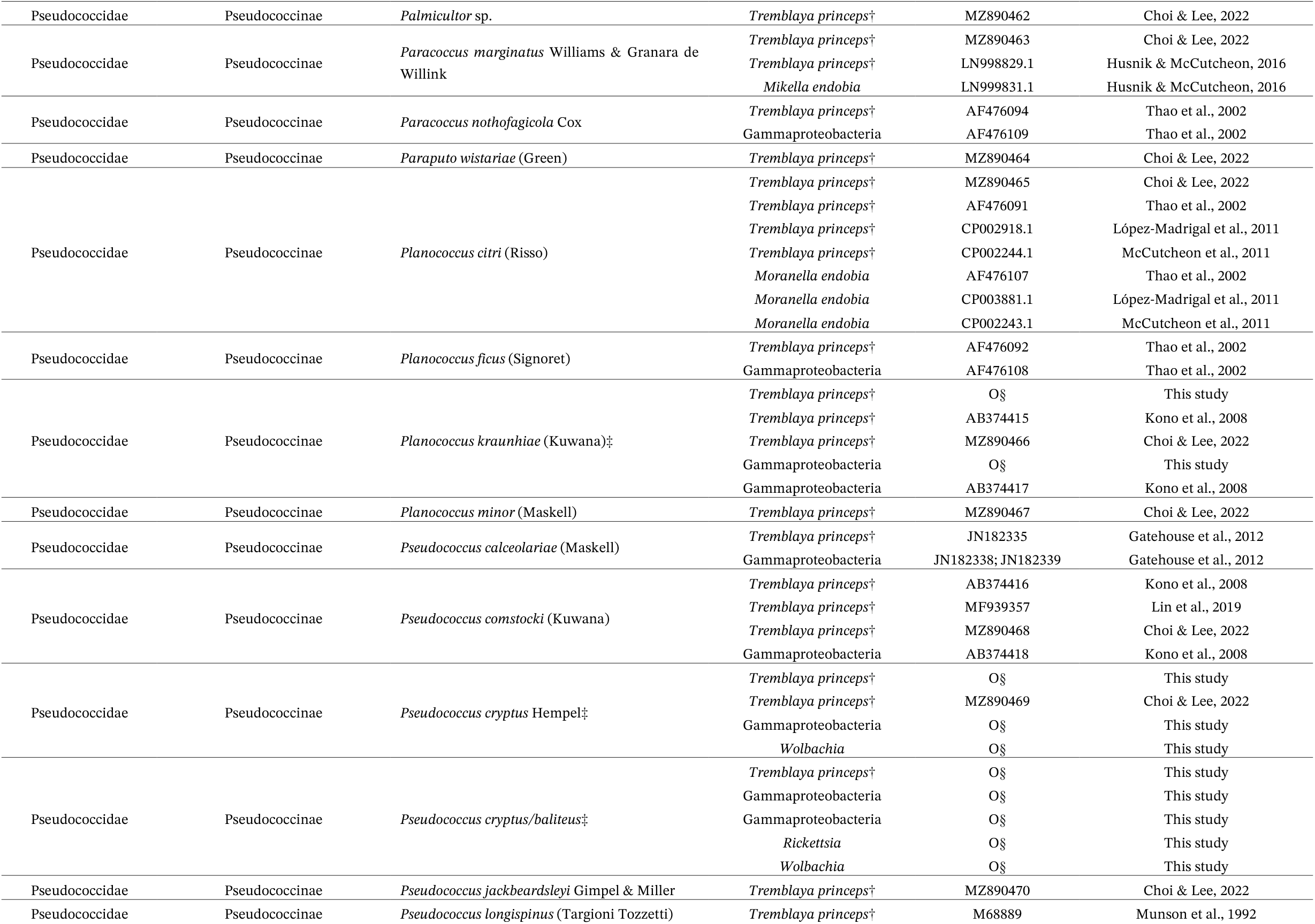
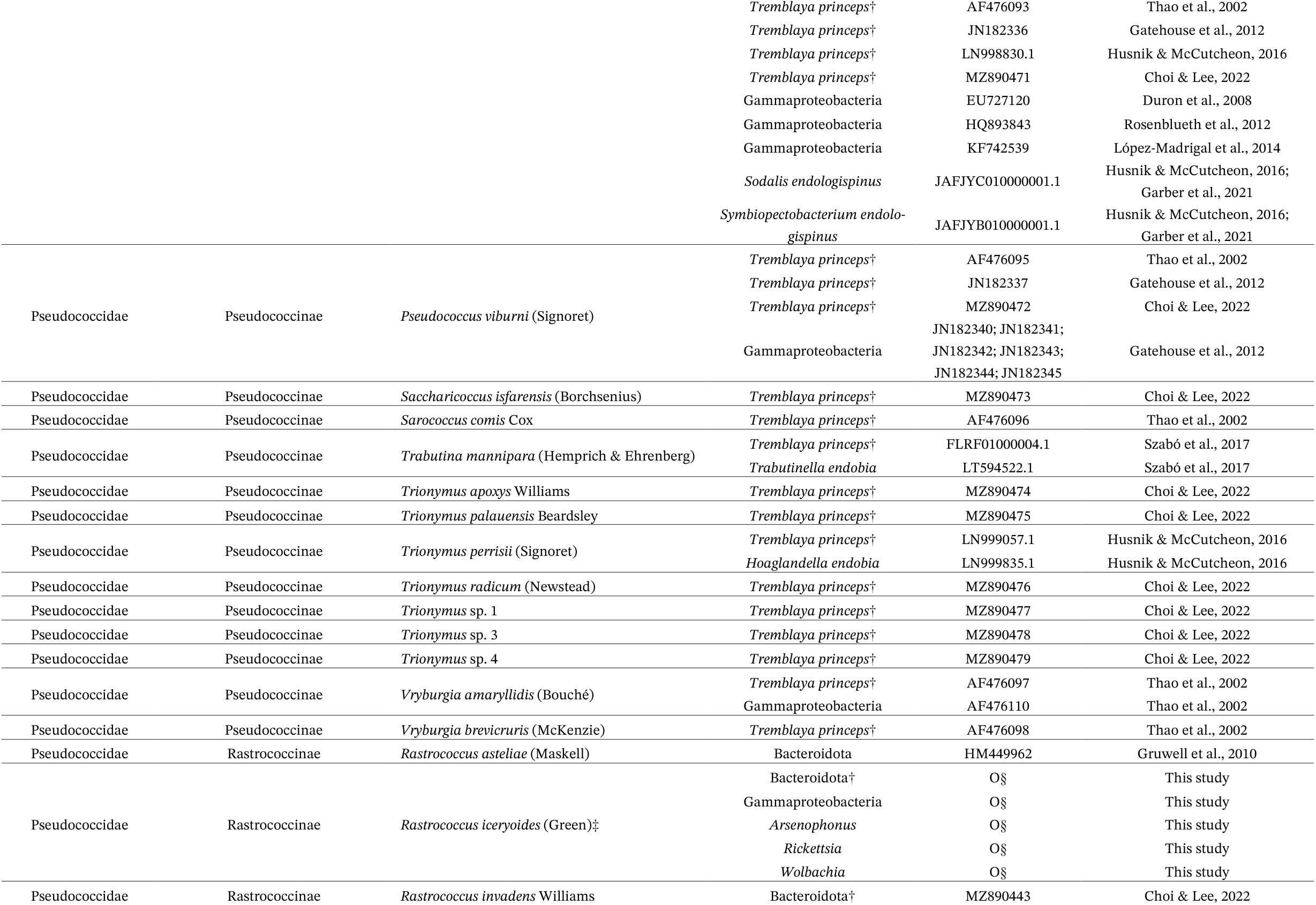
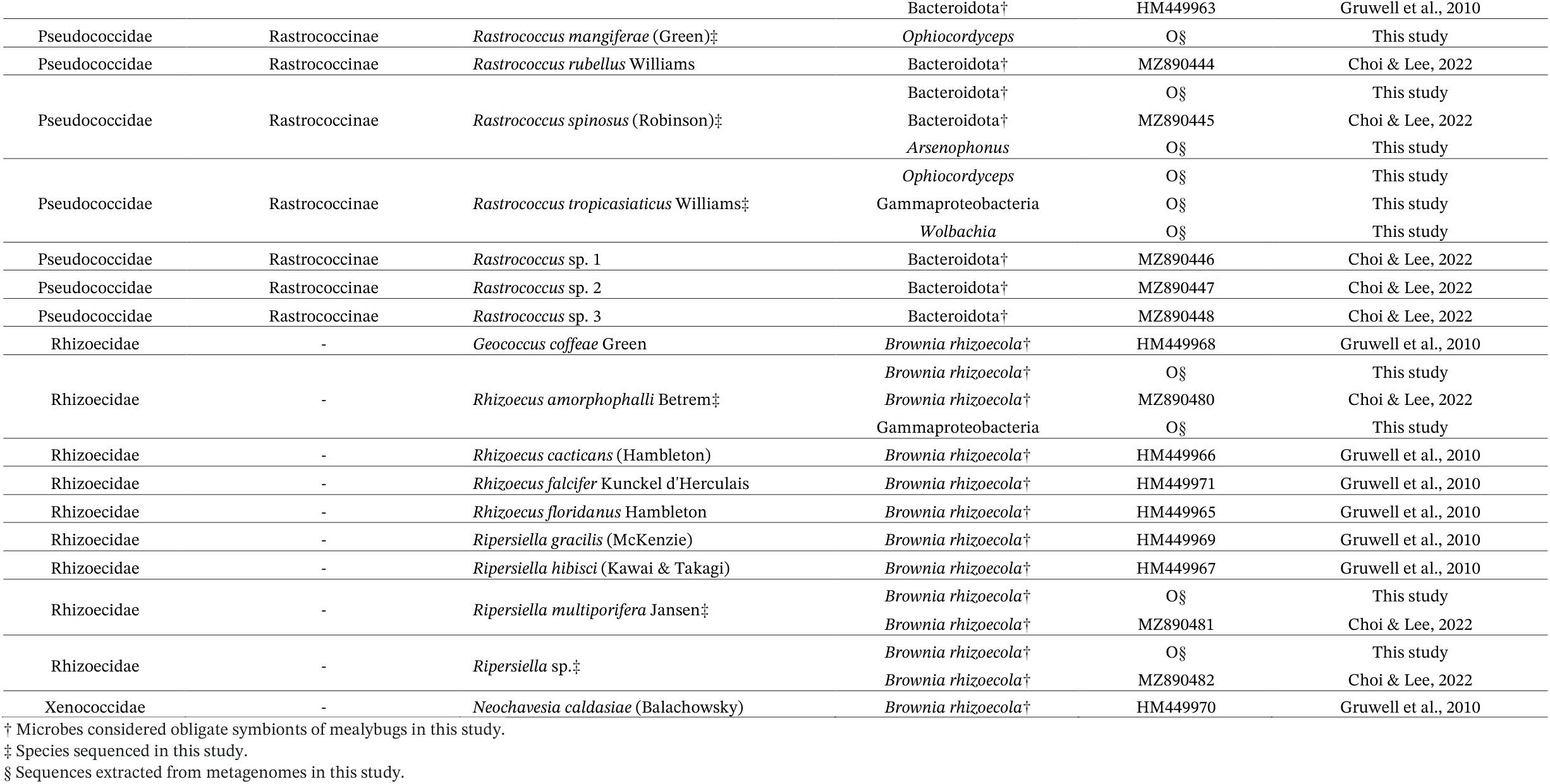
Symbionts of mealybugs screened from metagenome sequencing of this study and literatures. GenBank data include rRNA sequences and genomes of symbionts.

**Table S3.**
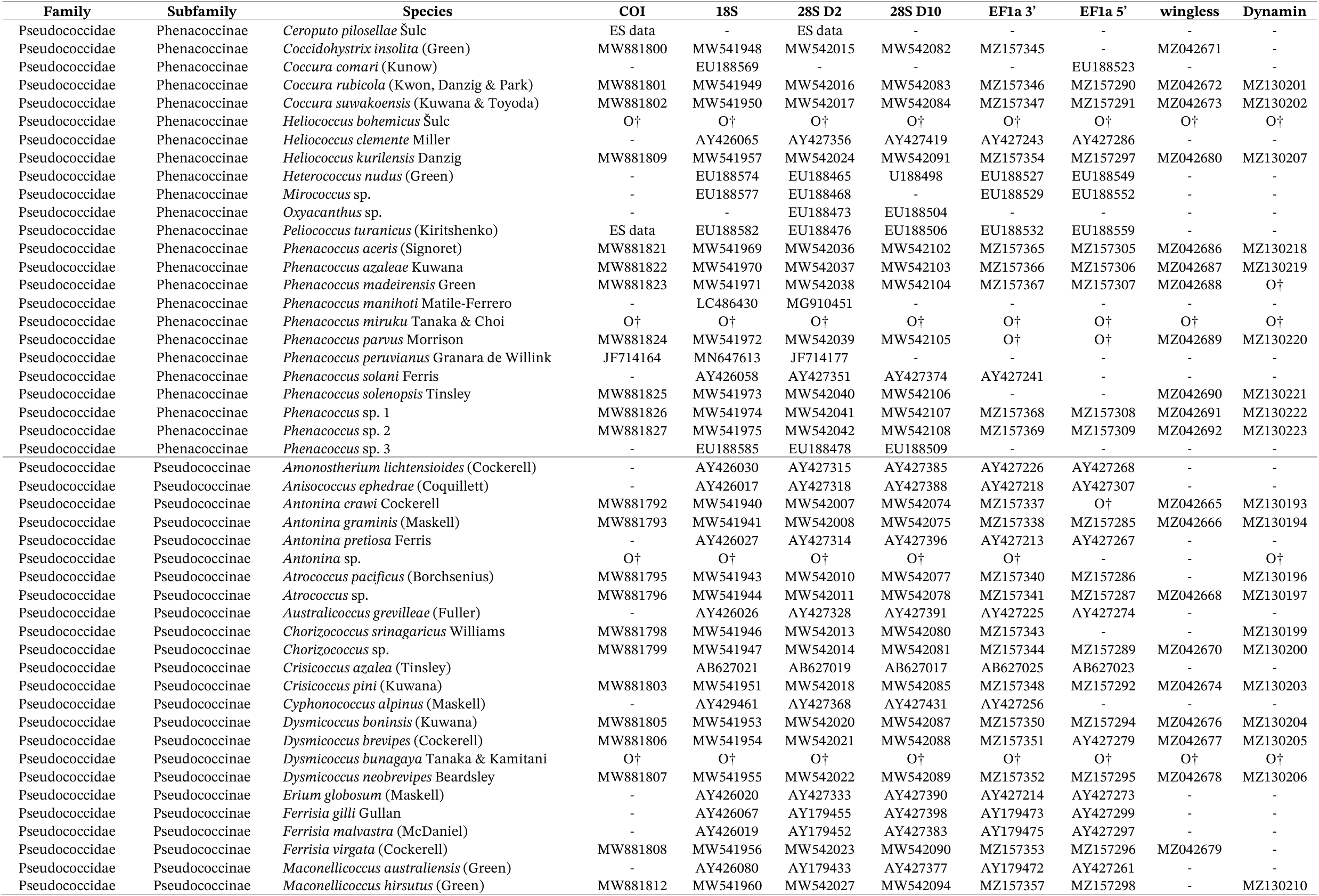
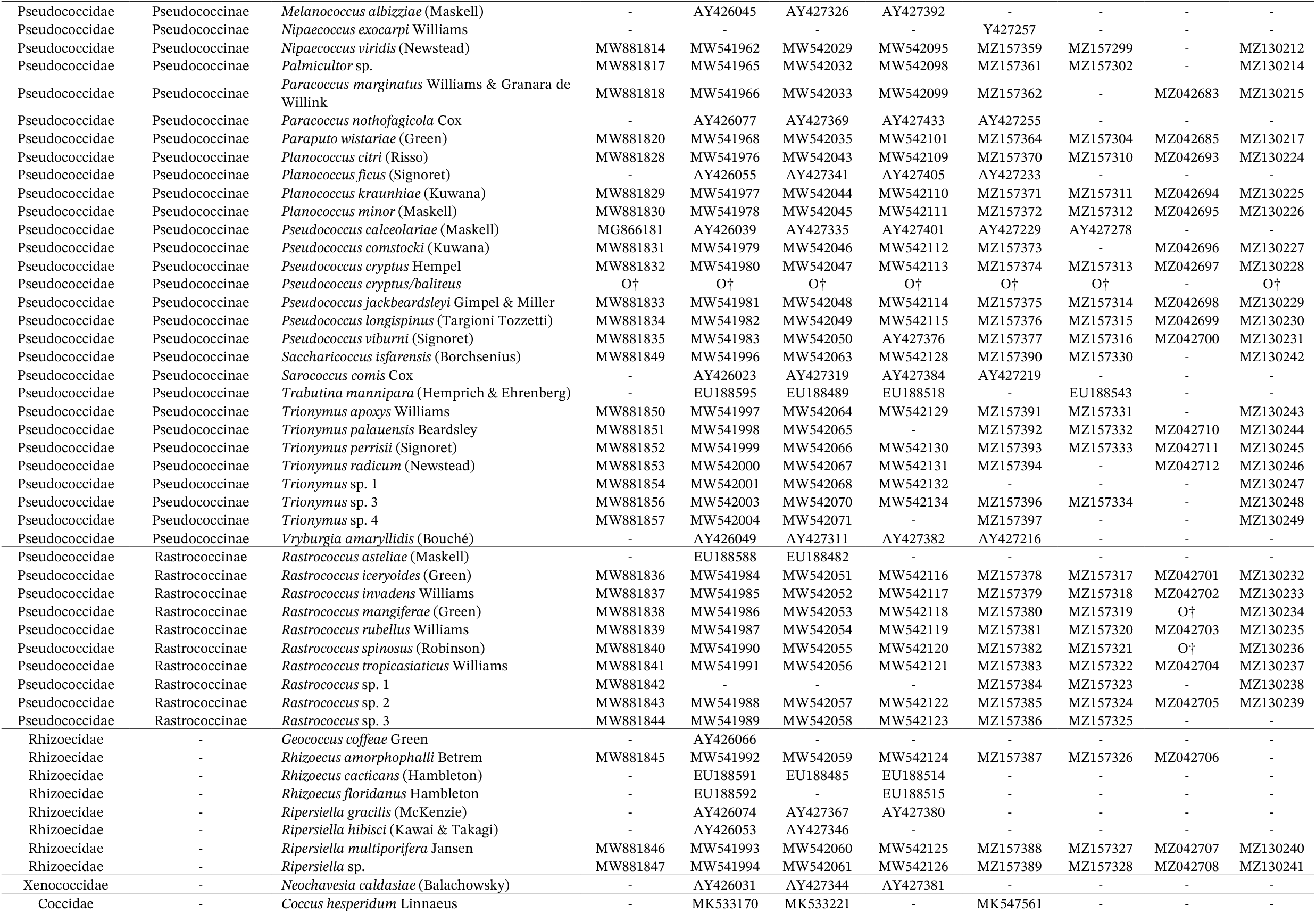
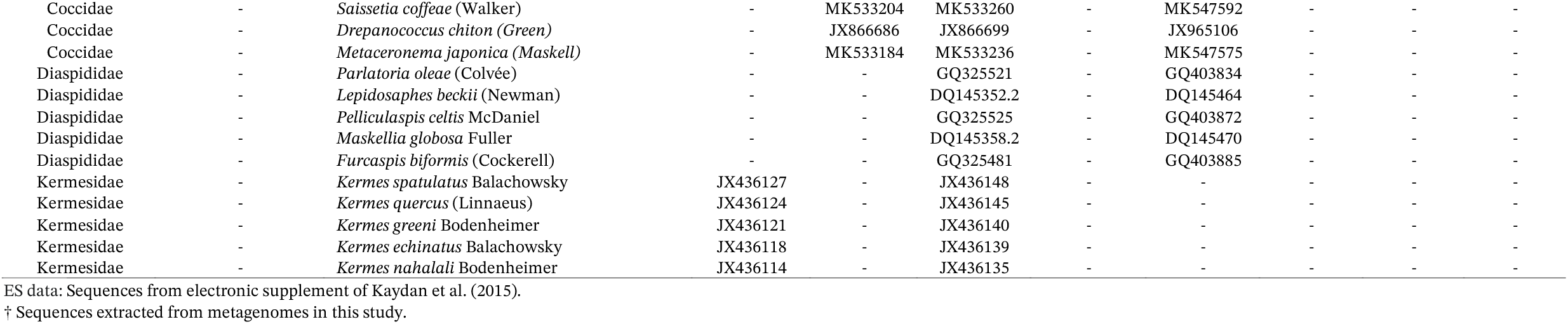
Taxa used for phylogenetic analysis of mealybugs with GenBank accession numbers.

## References

Agosta, S. J., Janz, N., & Brooks, D. R. (2010). How specialists can be generalists: resolving the “parasite paradox” and implications for emerging infectious disease. Zoologia, 27, 151–162.

Andrews, S. (2010). FastQC: A Quality Control Tool for High Throughput Sequence Data [Online]. Available online at: http://www.bioinformatics.babra-ham.ac.uk/projects/fastqc/

Bai, J., Zuo, Z., DuanMu, H., Li, M., Tong, H., Mei, Y., … & Li, F. (2024). Endosymbiont Tremblaya phenacola influences the reproduction of cotton mealybugs by regulating the mechanistic target of rapamycin pathway. The ISME Journal, 18(1), wrae052.

Bennett, G. M., & Moran, N. A. (2015). Heritable symbiosis: the advantages and perils of an evolutionary rabbit hole. Proceedings of the National Academy of Sciences, 112(33), 10169–10176.

Bouckaert, R., Vaughan, T. G., Barido-Sottani, J., Duchêne, S., Fourment, M., Gavryushkina, A., … & Matschiner, M. (2019). BEAST 2.5: An advanced software platform for Bayesian evolutionary analysis. PLoS computational biology, 15(4), e1006650.

Buchner, P. (1965). Endosymbiosis of animals with plant microorganisms. Interscience Publishers, 1–909.

Castresana, J. (2000). Selection of conserved blocks from multiple alignments for their use in phylogenetic analysis. Molecular Biology and Evolution, 17, 540–552.

Cates, R.G. (1980). Feeding patterns of monophagous, oligophagous, and polyphagous insect herbivores: the effect of resource abundance and plant chemistry. Oecologia, 46, 22–31.

Chen, S., Zhou, Y., Chen, Y., & Gu, J. (2018). fastp: an ultrafast all-in-one FASTQ preprocessor. Bioinformatics, 34(17), i884–i890.

Choi, J., & Lee, S. (2022). Higher classification of mealybugs (Hemiptera: Coccomorpha) inferred from molecular phylogeny and their endosymbionts. Systematic Entomology, 47, 354–370.

Chong, R. A., & Moran, N. A. (2018). Evolutionary loss and replacement of Buchnera, the obligate endosymbiont of aphids. The ISME journal, 12(3), 898–908.

Cornwallis, C. K., van’t Padje, A., Ellers, J., Klein, M., Jackson, R., Kiers, E. T., … & Henry, L. M. (2023). Symbioses shape feeding niches and diversification across insects. Nature ecology & evolution, 7(7), 1022–1044.

Correa, C. C., & Ballard, J. W. O. (2016). Wolbachia associations with insects: winning or losing against a master manipulator. Frontiers in Ecology and Evolution, 3, 153.

Crane, P. R., Friis, E. M., & Pedersen, K. R. (1995). The origin and early diversification of angiosperms. Nature, 374(6517), 27–33.

Dixon, A. F. G. (1977). Aphid ecology: life cycles, polymorphism, and population regulation. Annual Review of Ecology and Systematics, 329–353.

Douglas, A.E. (2016). How multi-partner endosymbioses function. Nature Reviews Microbiology, 14(12), 731–743.

Drummond, A. J., Ho, S. Y., Phillips, M. J., & Rambaut, A. (2006). Relaxed phylogenetics and dating with confidence. PLoS Biol, 4(5), e88.

Engl, T., Eberl, N., Gorse, C., Krüger, T., Schmidt, T. H., Plarre, R., … & Kaltenpoth, M. (2018). Ancient symbiosis confers desiccation resistance to stored grain pest beetles. Molecular Ecology, 27(8), 2095–2108.

Forister, M. L., Novotny, V., Panorska, A. K., Baje, L., Basset, Y., Butterill, P. T., … & Dyer, L. A. (2015). The global distribution of diet breadth in insect herbivores. Proceedings of the National Academy of Sciences, 112(2), 442–447.

Futuyma, D.J., & Agrawal, A.A. (2009). Macroevolution and the biological diversity of plants and herbivores. Proceedings of the National Academy of Sciences, 106(43), 18054–18061.

Garber, A. I., Kupper, M., Laetsch, D. R., Weldon, S. R., Ladinsky, M. S., Bjorkman, P. J., & McCutcheon, J. P. (2021). The evolution of interdependence in a fourway mealybug symbiosis. Genome biology and evolution, 13(8), evab123.

García Morales, M., Denno, B. D., Miller, D. R., Miller, G. L., Ben-Dov, Y. & Hardy, N. B. (2024). ScaleNet: a literature-based model of scale insect biology and systematics. Database. http://scalenet.info

Gil, R., Vargas-Chavez, C., López-Madrigal, S., Santos-García, D., Latorre, A., & Moya, A. (2018). Tremblaya phenacola PPER: an evolutionary beta-gammaproteobacterium collage. The ISME journal, 12(1), 124–135.

Grimaldi, D., & Engel, M. S. (2005). Evolution of the Insects. Cambridge University Press, 1–755.

Gruber-Vodicka, H. R., Seah, B. K., & Pruesse, E. (2020). phyloFlash: rapid small-subunit rRNA profiling and targeted assembly from metagenomes. Msystems, 5(5), e00920–20.

Gruwell, M. E., Hardy, N. B., Gullan, P. J., & Dittmar, K. (2010). Evolutionary relationships among primary endosymbionts of the mealybug subfamily Phenacoccinae (Hemiptera: Coccoidea: Pseudococcidae). Applied and Environmental Microbiology, 76(22), 7521–7525.

Gullan, P. J. & Kosztarab, M. (1997). Adaptations in scale insects. Annual Review of Entomology, 42, 23–50.

Gullan, P. J., & Cook, L. G. (2007). Phylogeny and higher classification of the scale insects (Hemiptera: Sternorrhyncha: Coccoidea). Zootaxa, 1668(1), 413–425.

Hammer, T. J., & Bowers, M. D. (2015). Gut microbes may facilitate insect herbivory of chemically defended plants. Oecologia, 179(1), 1–14.

Hansen, A. K., & Moran, N. A. (2014). The impact of microbial symbionts on host plant utilization by herbivorous insects. Molecular ecology, 23(6), 1473–1496.

Hardy, N.B., Peterson, D. A., & Normark, B. B. (2015). Scale insect host ranges are broader in the tropics. Biology letters, 11(12), 20150924.

Hoang, D. T., Chernomor, O., Von Haeseler, A., Minh, B. Q., & Vinh, L. S. (2018). UFBoot2: improving the ultra-fast bootstrap approximation. Molecular biology and evolution, 35(2), 518–522.

Hodgson, C. J., & Hardy, N. B., (2013). The phylogeny of the superfamily Coccoidea (Hemiptera: S ternorrhyn-cha) based on the morphology of extant and extinct macropterous males. Systematic Entomology, 38(4), 794–804.

Hodgson, C.J. (2020). A review of neococcid scale insects (Hemiptera: Sternorrhyncha: Coccomorpha) based on the morphology of the adult males. Zootaxa, 4765, 1–264.

Husnik, F., & McCutcheon, J. P. (2016). Repeated replacement of an intrabacterial symbiont in the tripartite nested mealybug symbiosis. Proceedings of the National Academy of Sciences, 113(37), E5416– E5424.

Husnik, F., Nikoh, N., Koga, R., Ross, L., Duncan, R. P., Fujie, M., … & Von Dohlen, C. D. (2013). Horizontal gene transfer from diverse bacteria to an insect genome enables a tripartite nested mealybug symbiosis. Cell, 153(7), 1567–1578.

Janson, E. M., Stireman, J. O., Singer, M. S., & Abbot, P. (2008). Phytophagous insect–microbe mutualisms and adaptive evolutionary diversification. Evolution: International Journal of Organic Evolution, 62(5), 997–1012.

Janz, N., & Nylin, S. (2008). The oscillation hypothesis of host-plant range and speciation. Specialization, speciation, and radiation: the evolutionary biology of herbivorous insects, 2008, 203–215.

Kaech, H., & Vorburger, C. (2021). Horizontal transmission of the heritable protective endosymbiont Hamiltonella defensa depends on titre and haplotype. Frontiers in microbiology, 11, 3577.

Kalyaanamoorthy, S., Minh, B. Q., Wong, T. K., von Haeseler, A., & Jermiin, L. S. (2017). Modelfinder: fast model selection for accurate phylogenetic estimates. Nature methods, 14(6): 587.

Katoh, K., & Standley, D. M. (2013). MAFFT multiple sequence alignment software v. 7. improvements in performance and usability. Molecular Biology and Evolution, 30, 772–780.

Kearse, M., Moir, R., Wilson, A., Stones-Havas, S., Cheung, M., Sturrock, S., … & Drummond, A. (2012). Geneious Basic: an integrated and extendable desktop software platform for the organization and analysis of sequence data. Bioinformatics, 28(12), 1647–1649.

Kiefer, J.S.T., Bauer, E., Okude, G., Fukatsu, T., Kaltenpoth, M., & Engl, T. (2023). Cuticle supplementation and nitrogen recycling by a dual bacterial symbiosis in a family of xylophagous beetles. The ISME Journal, 17(7), 1029–1039.

Koga, R., Nikoh, N., Matsuura, Y., Meng, X. Y., & Fukatsu, T. (2013). Mealybugs with distinct endosymbiotic systems living on the same host plant. FEMS microbiology ecology, 83(1), 93–100.

Kondo, T., Gullan, P. J., & Williams, D. J. (2008). Coccidology. The study of scale insects (Hemiptera: Sternorrhyncha: Coccoidea). Ciencia y Tecnología Agropecuaria, 9, 55–61.

Kozár, F. (2004). Ortheziidae of the World. Hungarian Academy of Sciences, Plant Protection Institute, Budapest, 1–525.

Kozár, F. & Konczné Benedicty, Z. (2007). Rhizoecinae of the world. Budapest: Plant Protection Institute. Hungarian Academy of Sciences, 1–617.

López-Madrigal, S., Beltrà, A., Resurrección, S., Soto, A., Latorre, A., Moya, A., & Gil, R. (2014). Molecular evidence for ongoing complementarity and horizontal gene transfer in endosymbiotic systems of mealybugs. Frontiers in microbiology, 5, 449.

Majerus, M. E. N., & Hurst, G. D. D. (1997). Ladybirds as a model system for the study of male-killing symbionts. Entomophaga, 42(1), 13–20.

McCutcheon, J. P., & Moran, N. A. (2007). Parallel genomic evolution and metabolic interdependence in an ancient symbiosis. Proceedings of the National Academy of Sciences, 104(49), 19392–19397.

McCutcheon, J. P., & Moran, N.A. (2010). Functional convergence in reduced genomes of bacterial symbionts spanning 200 My of evolution. Genome biology and evolution, 2, 708–718.

McCutcheon, J. P., & Moran, N.A. (2012). Extreme genome reduction in symbiotic bacteria. Nature Reviews Microbiology, 10(1), 13–26.

McCutcheon, J. P., & Von Dohlen, C.D. (2011). An interdependent metabolic patchwork in the nested symbiosis of mealybugs. Current Biology, 21(16), 1366–1372.

McCutcheon, J. P., & Moran, N. A. (2012). Extreme genome reduction in symbiotic bacteria. Nature Reviews Microbiology, 10(1), 13–26.

McKenzie, H.L. (1967). Mealybugs of California: with taxonomy, biology, and control of north American species (Homoptera, Coccoidea, Pseudococcidae). Berkeley, LA: University of California Press, 1–525.

Moran, N. A., Tran, P., & Gerardo, N. M. (2005). Symbiosis and insect diversification: an ancient symbiont of sap-feeding insects from the bacterial phylum Bacteroidetes. Appl. Environ. Microbiol., 71(12), 8802–8810.

Nguyen, L. T., Schmidt, H. A., from Haeseler, A., & Minh, B. Q. (2015). IQ-TREE: a fast and effective stochastic algorithm for estimating maximumlikelihood phylogenies. Molecular Biology and Evolution, 32, 268–274.

Normark, B. B., & Johnson, N. A. (2011). Niche explosion. Genetica, 139(5), 551–564.

Pagel, M., & Meade, A. (2006). Bayesian analysis of correlated evolution of discrete characters by reversi-ble-jump Markov chain Monte Carlo. The American Naturalist, 167(6), 808–825.

Pagel, M., Meade, A., & Barker, D. (2004). Bayesian estimation of ancestral character states on phylogenies. Systematic biology, 53(5), 673–684.

Percy, D. M., Page, R. D., & Cronk, Q. C. (2004). Plant–insect interactions: double-dating associated insect and plant lineages reveals asynchronous radiations. Systematic Biology, 53(1), 120–127.

Perlman, S. J., Hunter, M. S., & Zchori-Fein, E. (2006). The emerging diversity of Rickettsia. Proceedings of the Royal Society B: Biological Sciences, 273(1598), 2097–2106.

Peris, D., & Condamine, F. L. (2024). The angiosperm radiation played a dual role in the diversification of insects and insect pollinators. Nature Communications, 15(1), 552.

Prjibelski, A., Antipov, D., Meleshko, D., Lapidus, A., & Korobeynikov, A. (2020). Using SPAdes de novo assembler. Current protocols in bioinformatics, 70(1), e102.

Rambaut, A. (2009) FigTree, a graphical viewer of phylogenetic trees. Institute of Evolutionary Biology University of Edinburgh. http://tree.bio.ed.ac.uk/soft-ware/figtree.

Rambaut, A., Drummond, A.J., Xie, D., Baele, G, & Suchard, M.A. (2018). Posterior summarisation in Bayesian phylogenetics using Tracer 1.7. Systematic Biology, syy032.

Rosas-Pérez, T., Rosenblueth, M., Rincón-Rosales, R., Mora, J., & Martínez-Romero, E. (2014). Genome sequence of “Candidatus Walczuchella monophlebidarum” the flavobacterial endosymbiont of Llaveia axin axin (Hemiptera: Coccoidea: Monophlebidae). Genome biology and evolution, 6(3), 714–726.

Sabree, Z. L., Kambhampati, S., & Moran, N. A. (2009). Nitrogen recycling and nutritional provisioning by Blattabacterium, the cockroach endosymbiont. Proceedings of the National Academy of Sciences, 106(46), 19521–19526.

Sabree, Z. L., Huang, C. Y., Okusu, A., Moran, N. A., & Normark, B. B. (2013). The nutrient supplying capabilities of Uzinura, an endosymbiont of armoured scale insects. Environmental Microbiology, 15(7), 1988–1999.

Santos-Garcia, D., Juravel, K., Freilich, S., Zchori-Fein, E., Latorre, A., Moya, A., … & Silva, F. J. (2018). To B or not to B: comparative genomics suggests Ar-senophonus as a source of B vitamins in whiteflies. Frontiers in Microbiology, 9, 2254.

Slove, J., & Janz, N. (2011). The relationship between diet breadth and geographic range size in the butterfly subfamily Nymphalinae—a study of global scale. PLoS ONE, 6, e16057.

Strong, D.R., Lawton, J.H., & Southwood, R. (1984). Insects on Plants. Cambridge, MA: Harvard Univ. Press, Cambridge, 1–320.

Sudakaran, S., Retz, F., Kikuchi, Y., Kost, C., & Kaltenpoth, M. (2015). Evolutionary transition in symbiotic syndromes enabled diversification of phytophagous insects on an imbalanced diet. The ISME journal, 9(12), 2587–2604.

Sudakaran, S., Kost, C., & Kaltenpoth, M. (2017). Symbiont acquisition and replacement as a source of ecological innovation. Trends in Microbiology, 25(5), 375–390.

Szklarzewicz, T., Kalandyk-Kołodziejczyk, M., & Michalik, A. (2022). Ovary structure and symbiotic associates of a ground mealybug, Rhizoecus albidus (Hemip-tera, Coccomorpha: Rhizoecidae) and their phylogenetic implications. Journal of Anatomy, 241(3), 860–872.

Talavera, G. & Castresana, J. (2007). Improvement of phylogenies after removing divergent and ambiguously aligned blocks from protein sequence alignments. Systematic Biology, 56(4), 564–577.

Thao, M. L., Gullan, P. J., & Baumann, P. (2002). Secondary (γ-Proteobacteria) endosymbionts infect the primary (β-Proteobacteria) endosymbionts of mealybugs multiple times and coevolve with their hosts. Applied and Environmental Microbiology, 68(7), 3190–3197.

Tsai, C. W., Rowhani, A., Golino, D. A., Daane, K. M., & Almeida, R. P. (2010). Mealybug transmission of grapevine leafroll viruses: an analysis of virus–vector specificity. Phytopathology, 100, 830–834.

Urban, J. M., & Cryan, J. R. (2012). Two ancient bacterial endosymbionts have coevolved with the planthoppers (Insecta: Hemiptera: Fulgoroidea). BMC evolutionary biology, 12(1), 87.

Vaidya, G., Lohman, D. J., & Meier, R. (2011). SequenceMatrix: concatenation software for the fast assembly of multi-gene datasets with character set and codon information. Cladistics, 27, 171–180.

Vea, I. M., & Grimaldi, D. A. (2015). Diverse new scale insects (Hemiptera: Coccoidea) in amber from the Cretaceous and Eocene with a phylogenetic framework for fossil Coccoidea. American Museum Novitates, 1–80.

Vea, I. M., & Grimaldi, D. A. (2016). Putting scales into evolutionary time: the divergence of major scale insect lineages (Hemiptera) predates the radiation of modern angiosperm hosts. Scientific Reports, 6, 23487.

von Dohlen, C. D., Kohler, S., Alsop, S. T., & McManus, W. R. (2001). Mealybug β-proteobacterial endosymbionts contain γ-proteobacterial symbionts. Nature, 412(6845), 433–436.

Vorburger, C., Gehrer, L., & Rodriguez, P. (2010). A strain of the bacterial symbiont Regiella insecticola protects aphids against parasitoids. Biology letters, 6(1), 109–111.

Vorburger, C. (2021). Defensive Symbionts and the Evolution of Parasitoid Host Specialization. Annual review of entomology, 67, 329–346.

Werren, J. H., Baldo, L., & Clark, M. E. (2008). Wolbachia: master manipulators of invertebrate biology. Nat. Rev. Microbiol., 6, 741–751.

Williams, D. J. (2004b) Mealybugs of southern Asia. Kuala Lumpur: Natural History Museum Jointly with Southdene, 1–896.

Winkler, I. S., & Mitter, C. (2008). The phylogenetic dimension of insect-plant interactions: a review of recent evidence. Specialization, speciation, and radiation: the evolutionary biology of herbivorous insects, 240–263.

## Supplementary references

Duron, O., Bouchon, D., Boutin, S., Bellamy, L., Zhou, L., Engelstädter, J., & Hurst, G. D. (2008). The diversity of reproductive parasites among arthropods: Wolbachia do not walk alone. BMC biology, 6(1), 27.

Garber, A. I., Kupper, M., Laetsch, D. R., Weldon, S. R., Ladinsky, M. S., Bjorkman, P. J., & McCutcheon, J. P. (2021). The evolution of interdependence in a four-way mealybug symbiosis.

Gatehouse, L. N., Sutherland, P., Forgie, S. A., Kaji, R., & Christeller, J. T. (2012). Molecular and histological characterization of primary (Betaproteobacteria) and secondary (Gammaproteobacteria) endosymbionts of three mealybug species. Appl. Environ. Microbiol., 78(4), 1187–1197.

Gil, R., Vargas-Chavez, C., López-Madrigal, S., Santos-García, D., Latorre, A., & Moya, A. (2018). Tremblaya phenacola PPER: an evolutionary beta-gammaproteobacterium collage. The ISME journal, 12(1), 124–135.

Gruwell, M. E., Hardy, N. B., Gullan, P. J., & Dittmar, K. (2010). Evolutionary relationships among primary endosymbionts of the mealybug subfamily Phenacoccinae (Hemiptera: Coccoidea: Pseudococcidae). Applied and environmental microbiology, 76(22), 7521–7525.

Husnik, F., & McCutcheon, J. P. (2016). Repeated replacement of an intrabacterial symbiont in the tripartite nested mealybug symbiosis. Proceedings of the National Academy of Sciences, 113(37), E5416–E5427.

Husnik, F., Nikoh, N., Koga, R., Ross, L., Duncan, R. P., Fujie, M., … & McCutcheon, J. P. (2013). Horizontal gene transfer from diverse bacteria to an insect genome enables a tripartite nested mealybug symbiosis. Cell, 153(7), 1567–1578.

Koga, R., Nikoh, N., Matsuura, Y., Meng, X. Y., & Fukatsu, T. (2013). Mealybugs with distinct endosymbiotic systems living on the same host plant. FEMS microbiology ecology, 83(1), 93–100.

Kono, M., Koga, R., Shimada, M., & Fukatsu, T. (2008). Infection dynamics of coexisting beta-and gammaproteobacteria in the nested endosymbiotic system of mealybugs. Applied and environmental microbiology, 74(13), 4175–4184.

López-Madrigal, S., Beltrà, A., Resurrección, S., Soto, A., Latorre, A., Moya, A., & Gil, R. (2014). Molecular evidence for ongoing complementarity and horizontal gene transfer in endosymbiotic systems of mealybugs. Frontiers in microbiology, 5, 449.

López-Madrigal, S., Latorre, A., Porcar, M., Moya, A., & Gil, R. (2011). Complete genome sequence of “Candidatus Tremblaya princeps” strain PCVAL, an intriguing translational machine below the living-cell status.

McCutcheon, J. P., & Von Dohlen, C. D. (2011). An interdependent metabolic patchwork in the nested symbiosis of mealybugs. Current biology, 21(16), 1366–1372.

Michalik, A., Michalik, K., Grzywacz, B., Kalandyk-Kołodziejczyk, M., & Szklarzewicz, T. (2019). Molecular characterization, ultrastructure, and transovarial transmission of Tremblaya phenacola in six mealybugs of the Phenacoccinae subfamily (Insecta, Hemip-tera, Coccomorpha). Protoplasma, 1–12.

Munson et al., 1992. Phylogenetic relationships of the endosymbionts of mealybugs (Homoptera: Pseudococcidae) based on 16S rDNA sequences

Rosenblueth, M., Sayavedra, L., Sámano-Sánchez, H., Roth, A., & Martínez-Romero, E. (2012). Evolutionary relationships of flavobacterial and enterobacterial endosymbionts with their scale insect hosts (Hemiptera: Coccoidea). Journal of evolutionary biolog

Szabó, G., Schulz, F., Toenshoff, E. R., Volland, J. M., Finkel, O. M., Belkin, S., & Horn, M. (2017). Convergent patterns in the evolution of mealybug symbioses involving different intrabacterial symbionts. The ISME journal, 11(3), 715–726.

